# GTcomplex: Spatial indexing-powered search and alignment of macromolecular complexes

**DOI:** 10.64898/2025.12.15.694356

**Authors:** Mindaugas Margelevičius

## Abstract

Structural alignment of macromolecular complexes is essential for understanding their function and evolution, yet existing methods often rely on aligning individual chains before inferring complex-level correspondences, leading to inaccuracies and inefficiencies. Here we present GTcomplex, a novel algorithm that employs spatial indexing to perform holistic complex-level alignment, directly deriving chain assignments from optimal global superpositions. Benchmarking on diverse datasets—including protein complexes, viral capsids, and nucleic acid complexes—demonstrates that GTcomplex achieves state-of-the-art accuracy with substantial speed improvements over current methods. These advances enable scalable, accurate comparison of compositionally diverse and large assemblies, facilitating structural annotation, evolutionary studies, and multimeric structure prediction. GTcomplex is available as a user-friendly software package and as a web service supporting high-throughput searches.

## Introduction

Macromolecular complexes are crucial functional assemblies that govern the complexity and regulation of cellular processes [1]. A comprehensive understanding of their three-dimensional (3D) structures at atomic resolution is essential for elucidating the molecular mechanisms that underpin life and for developing strategies to modulate these processes.

Rapid technological progress and recent advances in artificial intelligence (AI) have revolutionized computational structural biology. These developments have enabled accurate prediction of both monomeric structures [2, 3] and multimeric assemblies [4–6], complementing experimental methods such as X-ray crystallography and cryogenic electron microscopy (cryo-EM). Together, predicted and experimentally determined structures provide invaluable insights into the conformation, dynamics, and functional states of biological assemblies. However, the explosive growth in structural data now demands robust and powerful computational methods for analysis.

The high structural conservation of macromolecular complexes [7] across diverse species [8] facilitates knowledge transfer, much as protein structure conservation enables functional annotation through sequence and structure similarity. Similar structures and conserved structural motifs not only shed light on shared molecular mechanisms but also support structure modeling and experimental structure determination [9]. As the volume of available data expands, the likelihood of identifying related complexes increases, accelerating discovery and creating a self-reinforcing cycle between structure prediction, comparison, and understanding.

Structural alignment is a fundamental technique for detecting structural similarity by establishing atomic correspondences between structures. It not only provides an optimal superposition of their 3D shapes but also yields quantitative similarity scores and enables the assessment of topological similarity and evolutionary relationships.

In contrast to the alignment of monomeric structures or individual chains, aligning macromolecular complexes requires mapping corresponding subunits (chains). As a result, complex-complex alignment involves determining a set of sub-alignments between matched chains, thereby extending the challenge of structural comparison from single molecules to entire assemblies.

As the demand for methods to compare complex structures has increased, several algorithmic strategies have emerged. QSalign [10] aligns entire complexes based on the chain order in the input file, infers chain correspondences from the resulting alignment, and then realigns the complexes accordingly. Because its dynamic programming algorithm enforces sequential order, this approach can produce suboptimal chain mappings and less accurate global alignments.

A typical strategy is to align all possible pairs of individual chains independently and then assign correspondences based on compatible orientations under superposition. This approach is used by MM-align [11], SCPC [12], VAST+ [13], US-align [14], and Foldseek-Multimer [15] (Foldseek-MM). Although similar in conceptual workflow, these methods differ in how they select optimal chain mappings. VAST+ and Foldseek-MM cluster transformation matrices representing similar spatial superpositions and retain those from the largest clusters. MM-align applies combinatorial optimization; and SCPC and US-align employ greedy algorithms. A greedy algorithm was also implemented in MICAN-SQ [16] and TopMatch [17], which align non-overlapping molecular segments irrespective of chain membership, without assigning chain correspondences. In a different vein, mTM-align2 [18] combines measures of overall molecular shape similarity with similarities between individual chains evaluated independently of their spatial arrangement.

Each method has strengths and limitations. Greedy algorithms, while generally fast, do not guarantee globally optimal solutions and may require multiple computationally intensive iterations to converge. Clustering-based approaches, on the other hand, can efficiently capture locally consistent superpositions but may fail to identify the globally optimal alignment because they do not evaluate overall structural similarity during clustering. (Our benchmarks highlight the conditions under which such methods can fail.)

To address these challenges, we introduce GTcomplex, a computational method for the search and alignment of macromolecular complexes based on spatial indexing. Rigorous large-scale benchmarking indicates that GTcomplex achieves, on average, (i) higher accuracy than US-align, one of the most accurate and widely used complex alignment tools, and (ii) comparable or, in some cases, >100x faster runtime performance than Foldseek-MM, currently the fastest method available. Beyond its speed and accuracy, GTcomplex is universal in scope, supporting both protein and nucleic acid assemblies—a property previously attributed only to US-align.

## Results

### GTcomplex algorithm

Before outlining the algorithm, it is useful to emphasize the one-to-one correspondence between structure superposition and structural alignment. A transformation matrix can be computed from a set of aligned residue or nucleotide pairs, and conversely, a structural alignment can be directly derived from a given superposition.

The central idea behind GTcomplex is that the overall similarity of two macromolecular complexes is best captured by identifying an optimal global superposition first, followed by determining the chain assignments implied by that superposition. This contrasts with the common workflow in complex alignment, where one first aligns all individual chains and then infers an overall complex-complex superposition from these chain-level alignments.

For homo-oligomers and distantly related complexes such as CRISPR-Cas complexes [19], optimal chain-level superpositions may not be uniquely determined without considering how they contribute to global complex similarity. Individual chains may match equally well to multiple chains, or a chain-level optimum may not correspond to the globally optimal complex-complex alignment.

Since evaluating large numbers of candidate complex-level alignments is computationally more demanding than aligning individual chains, the latter strategy has traditionally been adopted for efficiency. GTcomplex overcomes these computational limitations by leveraging the spatial indexing framework introduced in GTalign [20] for chain-level alignment.

Spatial indexing enables alignment derivation from a given superposition in O(1) time, a stark contrast to the quadratic complexity of dynamic programming conventionally used for this task. As a result, computing alignments for entire complexes is no more expensive than doing so for individual chains. Moreover, spatial-index-based alignments naturally produce chain assignments, whereas dynamic programming cannot handle non-sequential chain-to-chain correspondences.

GTcomplex exploits these advantages by first identifying the most promising superpositions, then deriving chain assignments from each, and finally refining the alignment associated with the optimal superposition.

An overview of the GTcomplex workflow is shown in Fig. 1a. The process begins with spatial indexing of both structures (not shown), which takes only a small fraction of the total runtime and is performed concurrently with data reading.

**Fig. 1.**
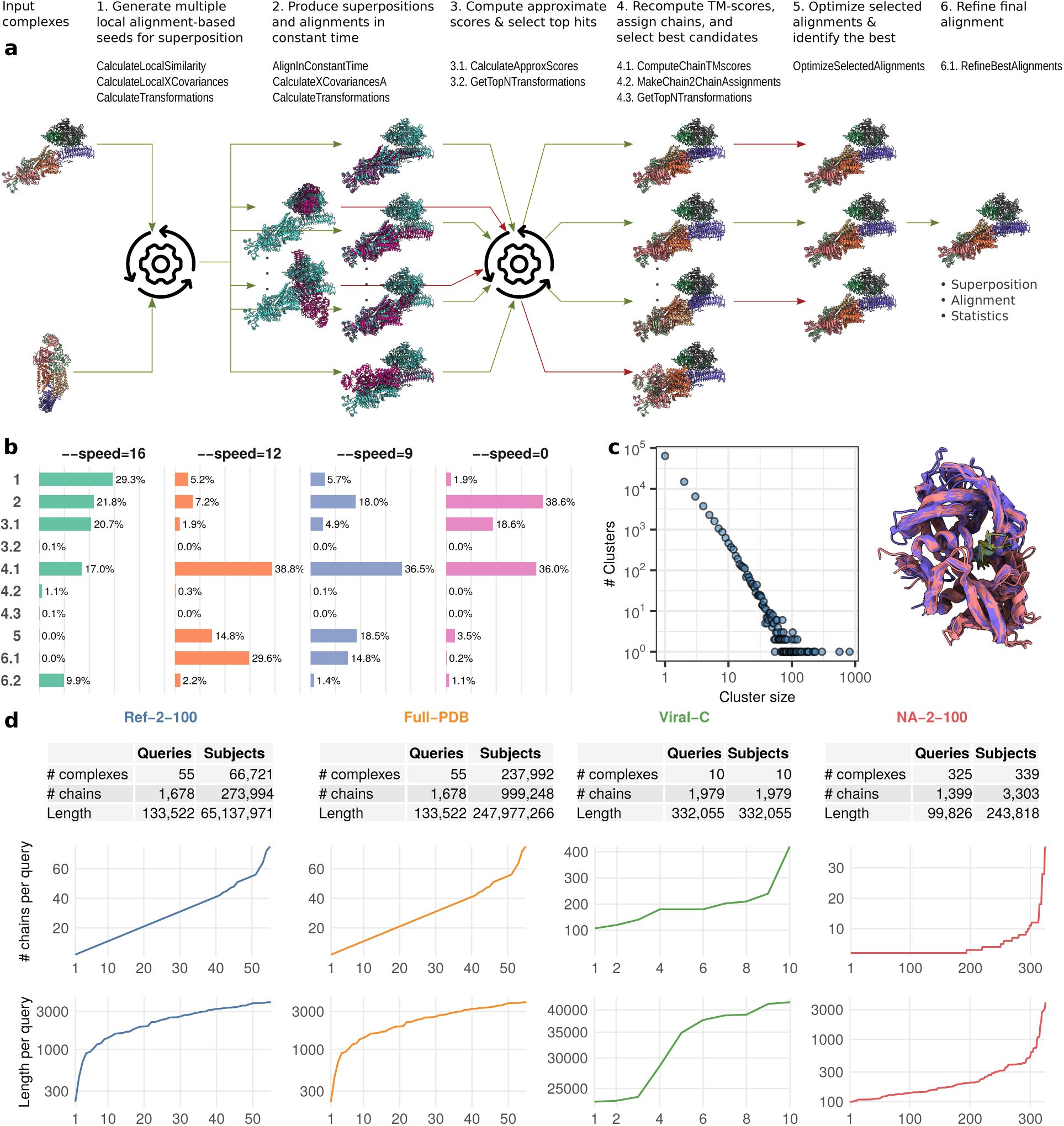
Overview of GTcomplex. a, The GTcomplex workflow comprises six main stages. In the figure, each stage is shown with a brief description, and the stage-specific algorithm names listed beneath each description correspond to those formally specified in Supplementary Section S2. Complexes are colored by chain except in Stage 2, where each complex is shown in a single distinct color to emphasize whole-complex processing. Red arrows indicate candidate superpositions that do not advance to the next stage. b, Runtime distribution (percentage of total runtime) across GTcomplex stages for different configurations specified by the --speed option. Stages 3, 4, and 6 are split into substages to highlight the computationally dominant components. Stage 6.2, although not listed explicitly in Panel a, applies the RefineBestAlignments procedure (associated with Stage 6.1) to perform fine-grained alignment refinement. c, Results of clustering the entire PDB using GTcomplex. The left panel shows the distribution of cluster sizes. The right panel shows the superimposition of all 214 members of the largest multi-chain cluster (primarily dimers), with structures colored by chain. d, Summary statistics for the Ref-2-100, Full-PDB, Viral-C, and NA-2-100 datasets used in benchmarking. Below each table, the distributions of chain counts and lengths (log scale) for the query complexes are shown, with each dataset represented in a distinct color. The x-axis indicates the index of queries sorted by chain count or by length, respectively.

The GTcomplex workflow proceeds through several main stages. Step 1: GTcomplex generates multiple local alignment seeds for initial superposition estimation, using local sequence similarity for nucleic acids or local structural similarity for proteins, with the seeds identified across whole complexes without prior chain decomposition. Because complexes exhibiting global similarity typically contain at least one locally similar region, these seeds provide starting points for estimating global superpositions and identifying globally consistent alignments. Step 2: From each initial superposition, alignments that may span multiple chains are derived using spatial indices, treating the entire complexes holistically (as indicated by uniform coloring in Fig. 1a). Step 3: These alignments are evaluated while preserving sequence order within their constituent chain-level alignments, consistent with the computation of TM-scores [21]. A predefined number of top-scoring alignments then advance to the next stage.

Step 4. For each superposition corresponding to a selected complex-level alignment, all chain pairs are realigned to compute chain-specific TM-scores that are consistent with the global (complex-level) superposition. These scores are then used to infer chain assignments, and the resulting complex-level TM-scores—computed as the sum of TM-scores across the assigned chain pairs—are used to select the highest-scoring superpositions for further analysis.

Step 5. These candidates are then optimized without altering chain assignments. Superpositions are refined using fragments of the full alignments, and TM-scores are recalculated.

Step 6. Finally, the candidate with the highest TM-score undergoes full refinement: chain-specific TM-scores are recomputed, chain assignments are updated, the alignment is adjusted, and the superposition is optimized as in Step 5.

GTcomplex is fully parallelized and highly efficient, enabling accurate complex alignments at scale.

### Runtime distribution across computational stages

The depth of analysis performed within each GTcomplex stage (Fig. 1a) is controlled by the --speed option. The fastest configuration, --speed=16, restricts Stage 2 to analyzing only highly locally similar alignments and retains only the top two candidates in Stage 3 for accurate TM-score recalculation in Stage 4.

Lowering the speed value increases the depth of exploration. For example, --speed=12 relaxes the local-similarity threshold in Stage 2, producing a larger set of superpositions for analysis, and increases the number of candidates advanced from Stage 3 to 16. With --speed=9, this number rises to 32, and the number of alignments subjected to optimization in Stage 5 increases from 1 to 5. The number of refinement iterations performed in Stage 6 increases to 2.

The deepest exploration level is provided by --speed=0. This variant removes the local-similarity threshold in Stage 2, allowing all initial superpositions to be considered. The number of alignments optimized in Stage 5 increases to 16, while the number of candidates reevaluated in Stage 4 (32) and the number of refinement iterations in Stage 6 (2) remain the same as in --speed=9.

The runtime distribution across the GTcomplex stages is shown in Fig. 1b, based on experiments using the Ref-2-100 dataset (described in the next subsection).

Across speed settings, Stage 4 is consistently one of the dominant contributors to the total runtime. This is due to the computation of chain-specific TM-scores via dynamic programming, which, although highly optimized and parallelized [22], remains computationally more intensive than the spatial index-based alignment used in Stage 2. Chain assignment (4.2) and candidate selection (4.3) contribute negligible overhead.

Stage 2—the spatial index-driven derivation of alignments from numerous initial superpositions—also contributes substantially, but for a different reason: the number of alignments can reach thousands per complex pair, particularly when using --speed=0.

Additional trends are notable. Under the fastest setting, --speed=16, Stage 1 consumes the largest portion of the total runtime because it generates the full pool of alignment seeds, only a fraction of which are explored further. Stages 5 and 6.1 are skipped entirely in --speed=16, leaving only Stage 6.2, which performs fine-grained refinement based on the RefineBestAlignments procedure (6.1).

Taken together, these results show that spatial indexing enables such efficient exploration of the superposition space that it often requires less time than the optimization of only a few selected candidate alignments.

### Diverse datasets for comprehensive evaluation

To rigorously evaluate GTcomplex across a range of biologically relevant scenarios, we constructed four structurally diverse benchmark datasets spanning major application domains for complex-level alignment: clustered protein complexes, the full PDB [23], large viral capsids, and nucleic acid complexes.

Ref-2-100 (diverse protein complexes). The first dataset, Ref-2-100, was constructed by clustering the entire PDB at a TM-score threshold of 0.5 and a minimum coverage of 70% using GTcomplex (Fig. 1c). Cluster representatives with at least two chains and at least 100 residues were selected to form the dataset, resulting in a total of 66,721 complexes. To ensure balanced coverage across oligomeric states, we randomly sampled 55 query complexes—one per group for each observed chain count—to search the full Ref-2-100 dataset.

Full-PDB (all PDB structures). The same 55 queries were also used to search the Full-PDB dataset, which included all macromolecular structures in the PDB [23]. For evaluation (next subsection), only subjects with at least two chains were considered, as monomeric structures can be appropriately assessed using single-chain alignment tools and would dilute the complex-specific assessment. Compared with the intentionally diverse Ref-2-100 dataset, Full-PDB represents a realistic large-scale search scenario for complex alignment.

Viral-C (large viral assemblies). To assess performance on extremely large complexes and the ability to detect structural relationships between them, we compiled the Viral-C dataset, consisting of ten viral capsid assemblies.

NA-2-100 (nucleic acid complexes). To evaluate performance on nucleic acid complexes, the NA-2-100 dataset included all PDB nucleic acid entries with at least two chains and at least 100 nucleotides. Complexes shorter than 4,000 nucleotides were selected as queries.

Summary statistics for all datasets as well as the distributions of chain counts and lengths for the query complexes are shown in Fig. 1d.

We additionally constructed three benchmark datasets from PDB biological assemblies, which are described together with the corresponding benchmarking results below.

### Evaluation framework and accuracy metrics

We benchmarked GTcomplex against the most widely used and actively maintained complex-alignment tools: US-align [14]—the successor to MM-align [11]—and the recently introduced Foldseek-MM [15]. Other methods reviewed in the Introduction were not included due to unavailability, lack of chain assignment functionality, or design limitations (e.g., restriction to complexes with the same number of subunits). MM-align, whose runtime can be orders of magnitude slower than US-align, was excluded from the benchmarks.

For both US-align and Foldseek-MM, we evaluated two commonly used configurations: the regular and fast (-fast) variants of US-align and Foldseek-MM together with Foldseek-MM-TM, the latter using TM-align [24] to produce chain-level alignments. To examine how GTcomplex’s --speed parameter balances accuracy and computational cost, we included several GTcomplex configurations spanning the full range of exploration depth.

Accuracy metrics. Alignment accuracy was evaluated using three complementary metrics: TM-score, GDT TS (proteins), and RMSD.

TM-score [21] and GDT TS [25] quantify global structural similarity between superposed complexes and range from 0 to 1. Whereas the TM-score can be viewed as a length-normalized sum of terms inversely proportional to the squared distances between aligned atoms, GDT TS linearly combines the fractions of residues aligned within four distance thresholds (1, 2, 4, and 8 *) and is therefore more conservative than the TM-score for large complexes.

Both metrics can be normalized by the length of either the query or the subject complex. In the main benchmarks, we report scores normalized by the length of the shorter complex, enabling the detection of all significant matches, including cases where a smaller complex aligns to a region of a larger one. For completeness, the Supplementary Information also reports results normalized by the query length, a setting that favours matches to complexes of similar or greater length while downweighting smaller complexes that may nonetheless share high local similarity with the query.

TM-score and GDT TS do not penalize mismatched or poorly aligned regions. Consequently, divergent segments or even entire chains included in an alignment are not reflected in these scores. In contrast, RMSD quantifies the average atomic deviation within the aligned region and is therefore sensitive to alignment inaccuracies. To complement the more permissive global metrics, we also report RMSD.

### GTcomplex achieves state-of-the-art accuracy with high efficiency

Benchmarking across all four datasets (Fig. 2) identified GTcomplex as the most accurate complex-alignment method, as measured by TM-score, the objective optimized by all evaluated tools. For the most exhaustive configuration, GTcomplex --speed=0, the cumulative TM-scores of top hits exceed those of US-align on every dataset: 4,617.3 vs. 4,508.4 (Ref-2-100), 23,736.1 vs. 23,164.3 (Full-PDB), 24.5 vs. 24.2 (Viral-C), and 1,915.7 vs. 1,914.0 (NA-2-100). These improvements translate into 23% and 18% more significant matches (TM-score ≥ 0.5) on Ref-2-100 and Full-PDB, respectively (5,194 vs. 4,232 and 25,616 vs. 21,720). Both methods identify 29 high-scoring matches on Viral-C, and GTcomplex identifies 2,107 versus 2,104 matches for US-align on NA-2-100.

**Fig. 2.**
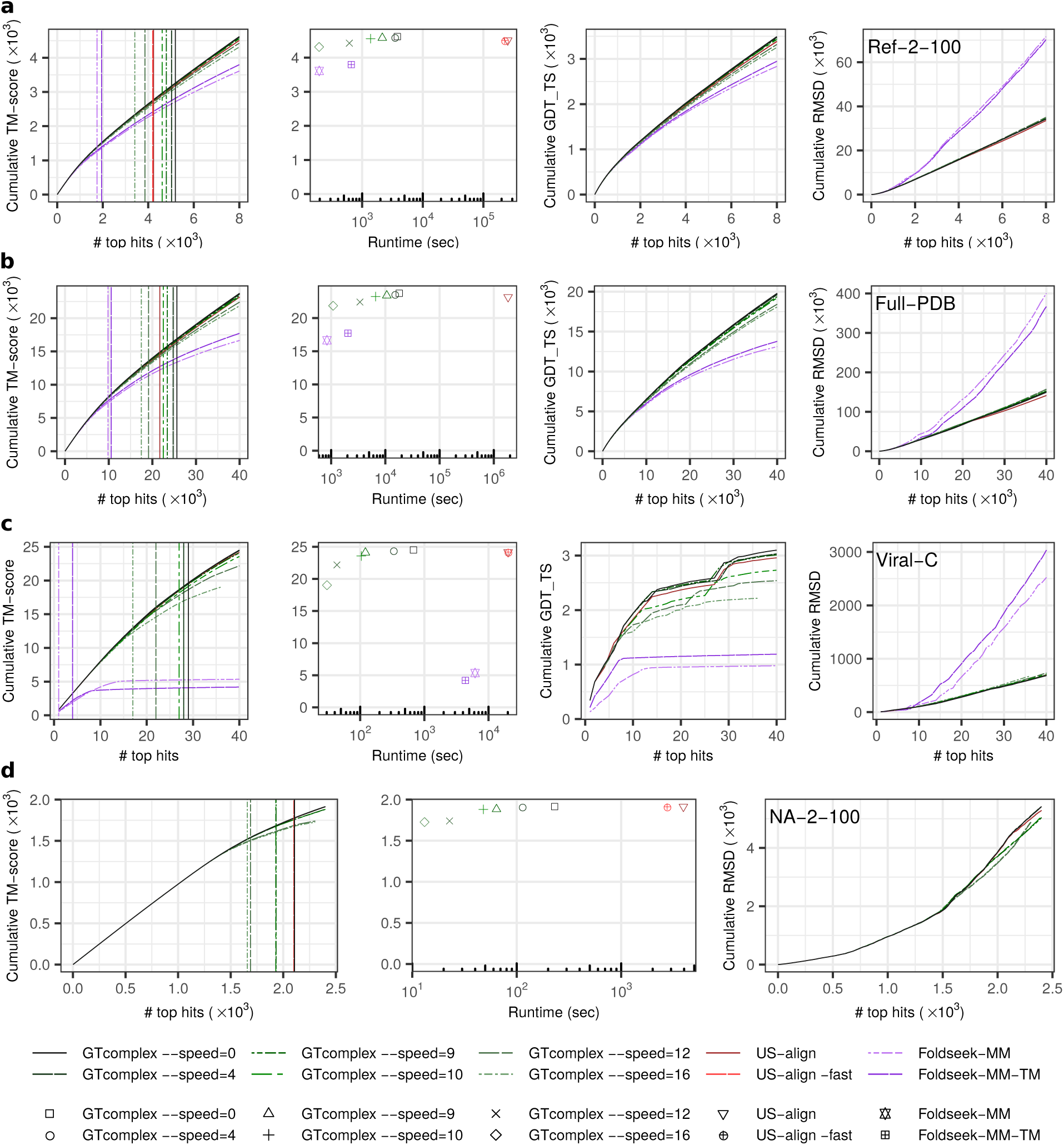
Benchmarking results on the Ref-2-100 (a), Full-PDB (b), Viral-C (c), and NA-2-100 (d) datasets. Left panels show cumulative TM-score as a function of the number of top-ranked alignments (ranked by TM-score). Vertical lines indicate the number of alignments with TM-score ≥ 0.5. Middle-left panels in (a–c) and the central panel in (d) plot cumulative TM-score versus runtime (seconds). Middle-right panels in (a–c) plot cumulative GDT_TS, and right panels plot cumulative RMSD, each as a function of the number of top-ranked alignments. Alignments are ranked by TM-score normalized by the length of the shorter complex (all evaluated tools optimize TM-score). In the left, middle-left (a–c), and central (d) panels, the TM-scores correspond to TM-align recalculations of the alignments (Methods), and the ranking of alignments is based on these recalculated TM-scores.

Because complex alignment methods may produce different alignments depending on whether the query is aligned to the subject or vice versa, we assessed the effect of retaining the higher-scoring of the two reciprocal alignments on the Ref-2-100 and NA-2-100 datasets. Although such a strategy is impractical for large-scale searches because it at least doubles the computational cost, it provides an upper bound on achievable alignment quality. The resulting scores differ only marginally from those obtained using a single alignment direction (Supplementary Fig. S1), leaving the conclusions of the large-scale benchmarks unchanged.

The advantage is also evident for the commonly identified top hits: TM-score scatter plots (Fig. 3) show that GTcomplex typically produces higher scores for the same hits. GDT TS results reinforce this trend, further demonstrating GTcomplex’s ability to produce accurate global alignments across a wide range of complex sizes and architectures (Fig. 2).

**Fig. 3.**
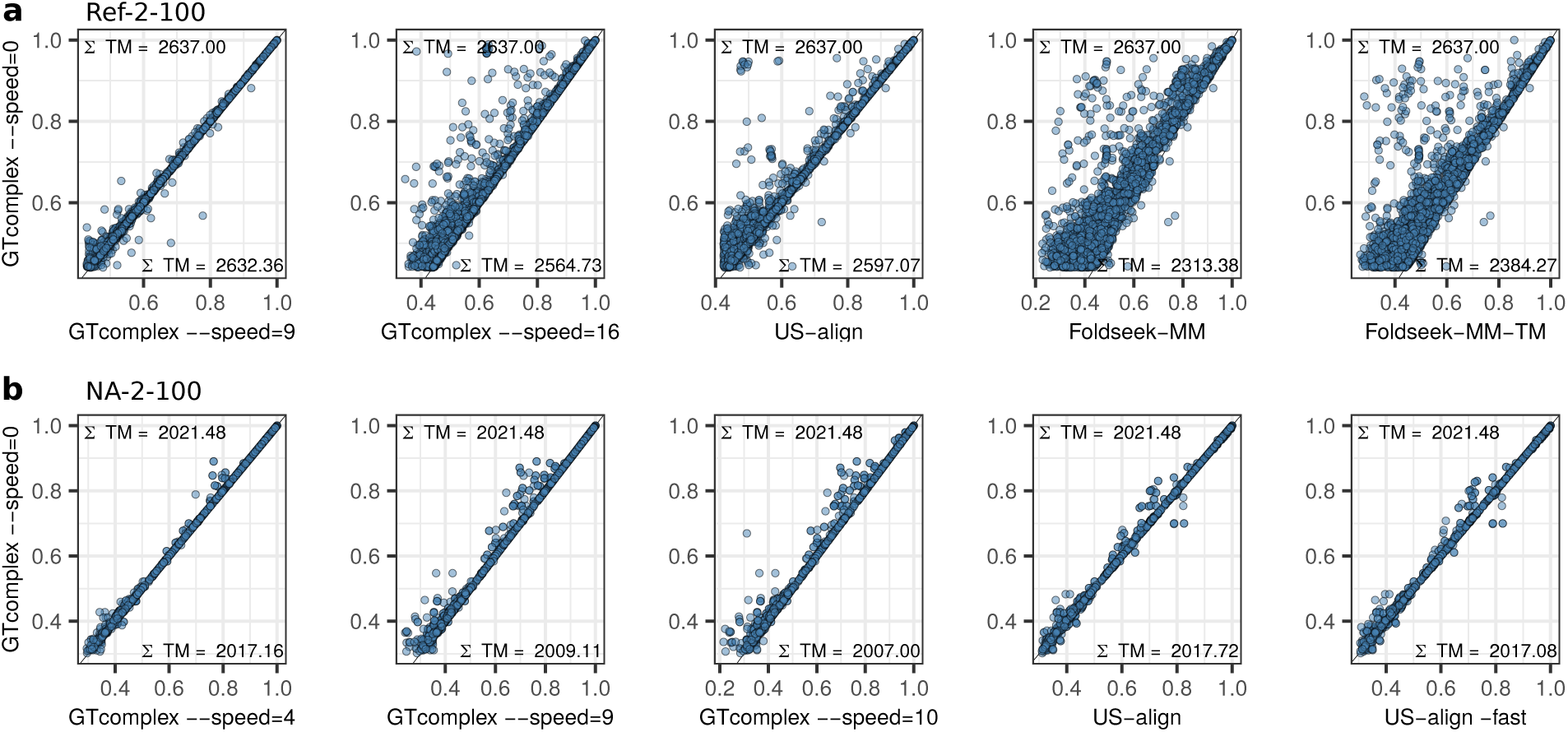
Head-to-head comparison of alignment accuracy across methods. Scatter plots compare TM-scores for the top 12,000 (a) and 4,800 (b) alignments produced by each tool, using only the alignments that all tools generated in common. Each point corresponds to an alignment between the same pair of complexes, with axes representing TM-scores from two different methods. Σ TM denotes the cumulative TM-score over all common alignments for each tool. a, Results for the Ref-2-100 dataset. b, Results for the NA-2-100 dataset.

GTcomplex exhibits slightly higher RMSDs than US-align on the Ref-2-100 and Full-PDB datasets. This effect arises because maximizing TM-score favors longer alignments with minimal local discrepancies, whereas extending an alignment naturally increases RMSD unless the two structures are nearly identical. Nonetheless, as shown in Fig. 2, the alignments produced by GTcomplex are consistently accompanied by higher TM-scores and GDT TS values, indicating that the method discovers true structural correspondences rather than introducing spurious mismatches.

These findings remain robust when evaluating alignments using TM-scores and GDT TS normalized by query length (Supplementary Fig. S2), a setting that favors matches to complexes of comparable or larger size and substantially reduces the number of possible matches with TM-score ≥ 0.5.

A key result is that GTcomplex achieves this accuracy at substantially lower computational cost. Even at its deepest search level (--speed=0), GTcomplex is about two orders of magnitude faster than US-align on the Ref-2-100 and Full-PDB datasets.

Moreover, GTcomplex offers a smooth accuracy-speed trade-off. More lightweight configurations recover most of the accuracy of --speed=0 while achieving great speedups. For example, GTcomplex --speed=16 completes the Ref-2-100 and Full-PDB searches in 195 s and 1,088 s, yielding speedups of 1,154x vs. US-align-fast (225,055 s) and 1,688x vs. US-align (1,836,471 s), respectively.

Speedups are smaller for Viral-C and NA-2-100. With runtimes of 30 s and 13 s, respectively, GTcomplex--speed=16 is 665x and 210x faster than US-align -fast (20,160 s and 2,750 s). This reduction in speedup reflects the smaller size of these datasets, as GTcomplex’s performance advantage becomes most pronounced in large-scale searches.

In comparison with Foldseek-MM, the fastest publicly available alternative, GTcomplex demonstrates competitive or superior efficiency. GTcomplex --speed=16 matches Foldseek-MM on Ref-2-100 (195 vs. 196 s), is moderately slower on Full-PDB (1,088 vs. 838 s), and is 206x faster on Viral-C (30 vs. 6,172 s).

Despite this efficiency, GTcomplex maintains substantially higher accuracy and sensitivity to significant structural similarity. GTcomplex --speed=16 identifies nearly twice as many matches with TM-score ≥ 0.5 as Foldseek-MM on Ref-2-100 and Full-PDB (3,413 vs. 1,752 and 17,464 vs. 9,831, respectively) and 17-fold more on Viral-C (17 vs. 1).

The most pronounced differences are observed in alignment accuracy, with RMSD highlighting these differences most clearly. RMSD analyses show that Foldseek-MM and Foldseek-MM-TM often produce alignments between spatially unrelated chains (Supplementary Fig. S3), an effect partially attributable to aligning individual chains without considering the context of full complexes.

### State-of-the-art performance extends to biological assemblies

The PDB contains both asymmetric units and biological assemblies. To evaluate GTcomplex exclusively on biologically relevant complexes, we constructed three additional datasets from PDB biological assemblies (Fig. 4a).

**Fig. 4.**
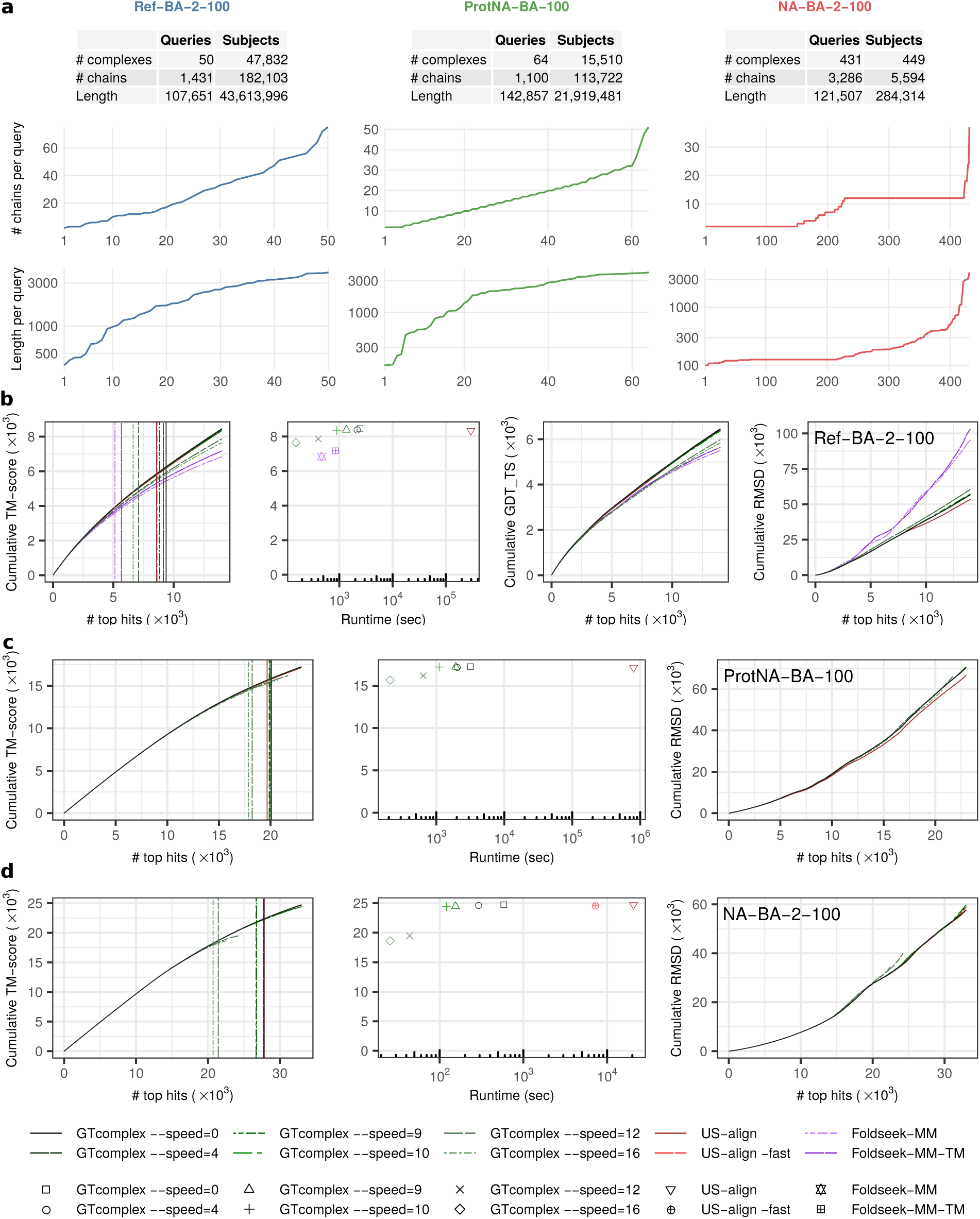
Benchmarking results on biological assembly datasets. a, Summary statistics for the Ref-BA-2-100, ProtNA-BA-100, and NA-BA-2-100 datasets. The statistics are represented as in Fig. 1d. b, Benchmarking results for Ref-BA-2-100. c, Benchmarking results for ProtNA-BA-100. d, Benchmarking results for NA-BA-2-100. Left panels show cumulative TM-score across top-ranked alignments ranked by TM-score. Vertical lines indicate the number of alignments with TM-score ≥ 0.5. The middle-left panel in (b) and the central panels in (c,d) show cumulative TM-score as a function of runtime (seconds). The middle-right panel in (b) plot cumulative GDT TS, and the right panels plot cumulative RMSD across top-ranked alignments. Alignments are ranked by TM-score normalized by the length of the shorter assembly. In the left panels, the middle-left panel of (b), and the central panel of (d), TM-scores correspond to TM-align recalculations of the alignments (Methods). For ProtNA-BA-100 (c), TM-scores are not recalculated.

Ref-BA-2-100 (diverse protein assemblies) was derived from the Ref-2-100 dataset by replacing structures with their corresponding biological assemblies. Assemblies containing at least two chains and 100–20,000 residues were included, with assemblies not exceeding 4,000 residues selected as queries.

ProtNA-BA-100 (protein-nucleic acid assemblies) comprised all biological assemblies containing both proteins and nucleic acids, with total lengths ranging from 100 to 10,000 residues. Query assemblies were represented by 64 randomly selected cluster representatives obtained using GTcomplex clustering at TM-score and coverage thresholds of 0.5 and 70%, respectively.

NA-BA-2-100 (nucleic acid assemblies) comprised all biological assemblies containing at least two chains and at least 100 nucleotides. Assemblies not exceeding 4,000 nucleotides were selected as queries.

Benchmarking on biological assemblies (Fig. 4) confirmed the superior accuracy of GTcomplex in terms of TM-score and GDT TS. For top hits, the cumulative TM-scores achieved by GTcomplex (--speed=0) and US-align are 8,454 vs. 8,339 for Ref-BA-2-100, 17,245 vs. 17,135 for ProtNA-BA-100, and 24,740 vs. 24,726 for NA-BA-2-100. For Ref-BA-2-100, the cumulative GDT TS is also higher for GTcomplex than for US-align (6,466 vs. 6,403). These conclusions are unchanged when for query-subject pairs the higher-scoring of the two reciprocal alignments was selected (Supplementary Fig. S4), indicating that score differences arising from alignment direction are relatively rare.

The same trend is observed for commonly identified top hits, for which GTcomplex generally produces higher TM-scores than other methods (Fig. 5).

**Fig. 5.**
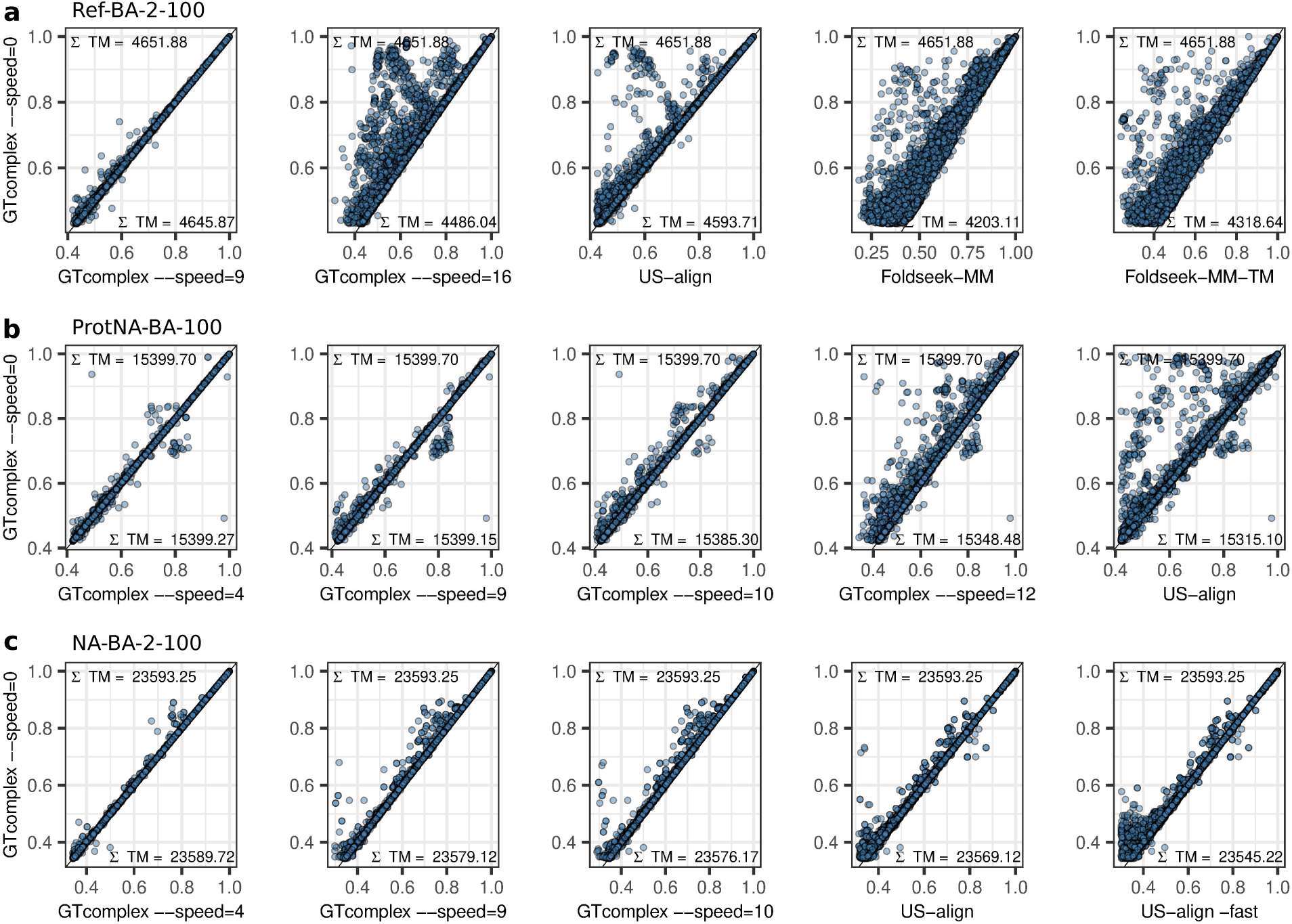
TM-score comparison for alignments identified by all methods. Scatter plots compare TM-scores for the top 18,000 (a), 25,000 (b), and 36,000 (c) alignments common to all methods. Each point represents the same query-subject assembly pair, with axes showing TM-scores from two methods. Σ TM denotes the cumulative TM-score across all common alignments for each method. a, Ref-BA-2-100. b, ProtNA-BA-100. c, NA-BA-

Consistent with the Ref-2-100 and Full-PDB benchmarks, the higher TM-scores obtained by GTcomplex on Ref-BA-2-100 and ProtNA-BA-100 are accompanied by slightly higher RMSDs than those produced by US-align. This behavior is expected because TM-score optimization may permit local deviations in favor of higher global structural similarity.

The conclusions are also robust to normalization of TM-score and GDT TS by query length (Supplementary Fig. S5). However, these results should be interpreted with caution because accurately aligned subject assemblies that are smaller than the query (Fig. 4b,c) are penalized in proportion to the query-subject length ratio.

Benchmarking on biological assemblies also demonstrates the high efficiency of GTcomplex (Fig. 4). The most exhaustive configuration (--speed=0) yields speedups of more than two orders of magnitude over US-align on the Ref-BA-2-100 and ProtNA-BA-100 datasets. The fastest configuration (--speed=16) preserves most of the accuracy while achieving speedups of 1,914x (154 s vs. 294,706 s) and 3,826x (209 s vs. 799,606 s), respectively. For the NA-BA-2-100 dataset, the smaller dataset size and lower nucleic-acid prefiltering threshold reduce speedups, yet GTcomplex still provides a 276x speedup over US-align -fast (26 s vs. 7,183 s). Compared with Foldseek-MM, GTcomplex --speed=16 is 3x faster on Ref-BA-2-100 (154 s vs. 466 s) while identifying substantially more significant matches with TM-score ≥ 0.5 (6,636 vs. 5,136).

### GTcomplex exhibits high sensitivity on a per-query basis

So far, the evaluations have been aggregated across all queries. Here, we assess sensitivity on a per-query basis. For each query, sensitivity is defined as the sum of TM-scores for up to five top-scoring alignments with TM-score ≥ 0.5, divided by five. Queries with fewer than five such alignments therefore receive lower sensitivity values that reflect both the number of confident matches identified and their average structural similarity. The fraction of queries exceeding a sensitivity threshold ranging between 0 and 1 thus characterizes a method’s ability to identify multiple confident matches and the average quality of those matches. The area under the resulting curve provides a summary measure of sensitivity across all queries, with a maximum value of 1 corresponding to five identical matches being identified for every query.

The results (Fig. 6) show that GTcomplex --speed=0 achieves the highest sensitivity among the evaluated methods across all datasets (the Viral-C dataset is omitted because it does not represent a large-scale search setting), with particularly pronounced improvements over the Foldseek-MM variants. The fastest GTcomplex configuration (--speed=16) also maintains high sensitivity, exceeding Foldseek-MM by 26%, 28%, and 31% on the Ref-2-100 (0.768 vs. 0.611), Full-PDB (0.859 vs. 0.672), and Ref-BA-2-100 (0.782 vs. 0.597) datasets, respectively.

**Fig. 6.**
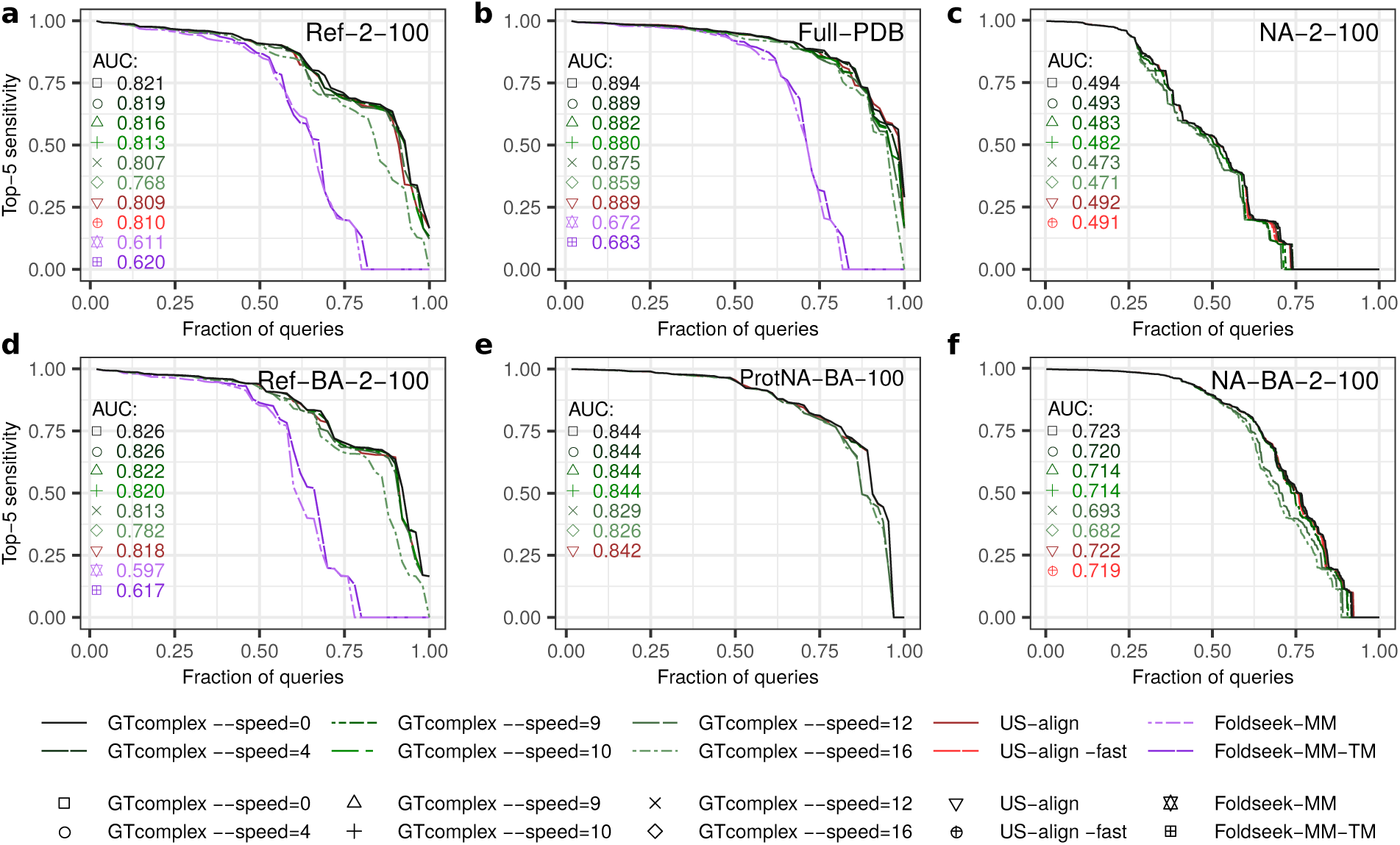
Per-query sensitivity analysis for the Ref-2-100 (a), Full-PDB (b), NA-2-100 (c), Ref-BA-2-100 (d), ProtNA-BA-100 (e), and NA-BA-2-100 (f) datasets. Curves show the fraction of queries exceeding a given sensitivity value. Sensitivity is defined as the sum of TM-scores for up to five top-scoring alignments with TM-score ≥ 0.5, divided by five, thereby rewarding both the number and quality of confident matches. AUC, area under the curve.

These results demonstrate that GTcomplex not only achieves high alignment accuracy but also consistently identifies multiple high-quality matches across diverse queries.

### Diverse biological examples demonstrate GTcomplex’s robustness

To illustrate GTcomplex’s robustness beyond benchmark statistics, we examine a set of diverse examples spanning protein complexes, nucleic acids, and large macromolecular assemblies.

The first example (Ref-BA-2-100) involves two functionally distinct complexes: the Fab fragment of the ABT007 antibody bound to the erythropoietin receptor (2jix) and the sDscam homodimer involved in cell-cell recognition (7y95). US-align and Foldseek-MM-TM align only one chain (Foldseek-MM-TM aligns the second chain highly inaccurately, as reflected by an RMSD of 17.4 *), whereas GTcomplex accurately aligns both chains of 7y95 (TM-score = 0.616), revealing a similar binding arrangement (Fig. 7a).

**Fig. 7.**
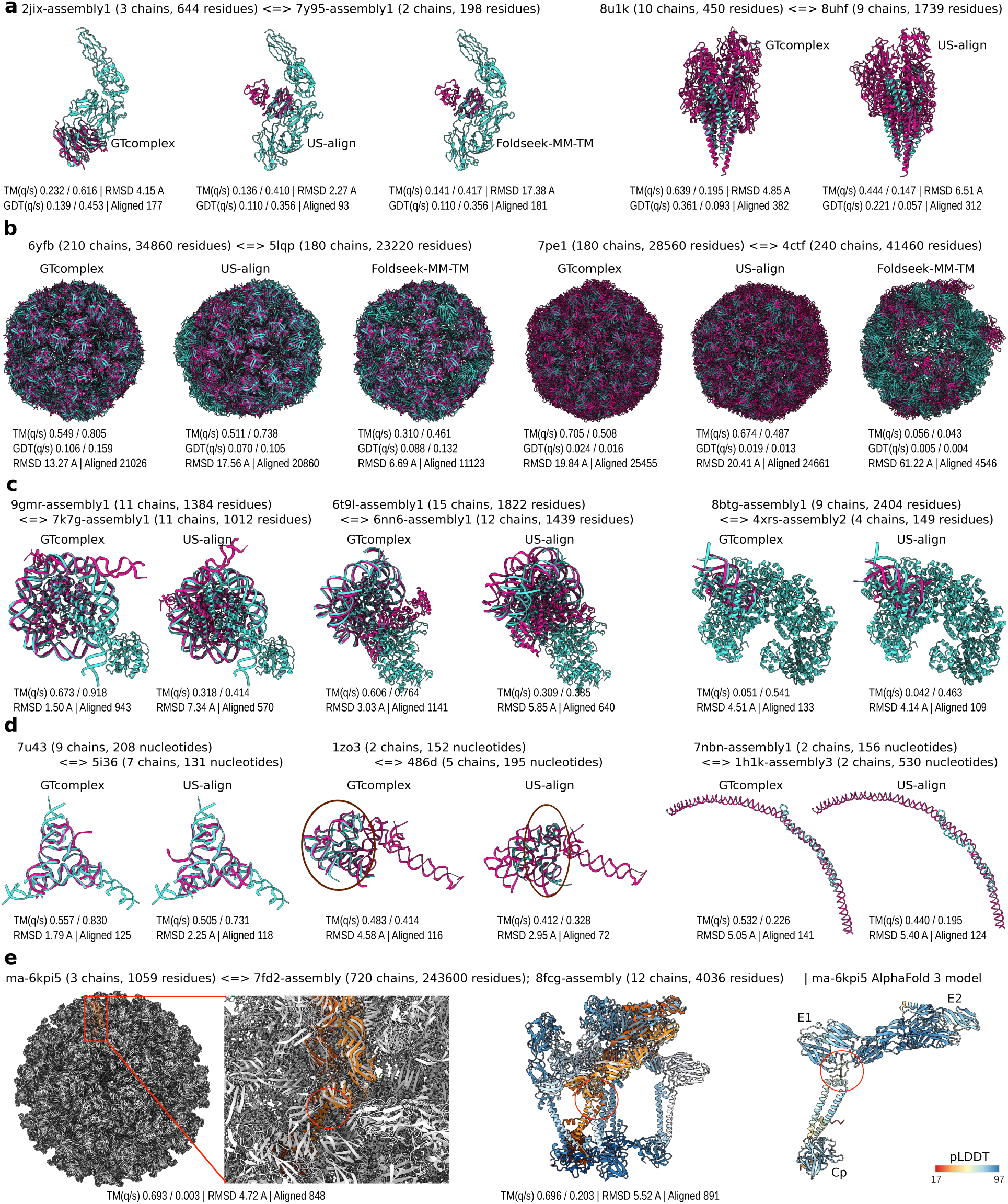
Illustrative examples demonstrating GTcomplex’s utility. a, Examples from benchmarking on the Ref-2-100 and Ref-BA-2-100 datasets. b, Examples from benchmarking on the Viral-C dataset. For Foldseek-MM-TM, only the chains included in the reported alignment are shown. c, Examples from benchmarking on the ProtNA-BA-100 dataset. d, Examples from benchmarking on the NA-2-100 and NA-BA-2-100 datasets. Oval frames highlight two chains aligned by GTcomplex compared with one chain aligned by US-align. e, Case study of structural model ma-6kpi5, a heterotrimer from Wenling crested flounder alphavirus. The AlphaFold 3 model (right) is colored by pLDDT confidence. Circular frames indicate low-confidence regions that cause the terminal α-helices of glycoproteins E1 and E2 to adopt orientations inconsistent with those observed in homologous viral complexes shown in the left and central panels. The ma-6kpi5 model is colored by chain using shades of yellow (except in the right panel), whereas the subject 8fcg is colored by chain using shades of blue. In panels a–d, query and subject complexes are shown in turquoise and red, respectively. Reported scores accompany each superposition. Normalization by query and subject lengths is denoted by q and s, respectively.

GTcomplex also confidently captures structural similarity between structurally diverse bacterial type IV pili family proteins [26] (Ref-2-100). Accurate chain mapping between the tight adhesion pilus (8u1k) and toxin-coregulated pilus (8uhf) structures from different bacteria enables GTcomplex to achieve a TM-score of 0.639 (Fig. 7a). In contrast, incorrect chain assignment by US-align yields a TM-score of 0.444, below the threshold typically considered indicative of confident structural similarity, whereas both Foldseek-MM variants fail to detect the relationship.

Examples from the Viral-C dataset further highlight GTcomplex’s advantage for large viral assemblies (Fig. 7b). For both a pair of single-stranded RNA bacteriophage capsids (6yfb and 5lqp) and the comparison of a brome mosaic virus capsomer assembly (7pe1) with the equine rhinitis A virus capsid (4ctf), GTcomplex achieves superior superpositions through more accurate chain assignments. Foldseek-MM-TM, in contrast, aligns only partial regions and often misorients subunits, leaving many chains mutually distant in space.

Examples from the ProtNA-BA-100 dataset demonstrate GTcomplex’s advantage for protein-nucleic acid assemblies (Fig. 7c). GTcomplex confidently identifies the evident relationship between a nucleosome in complex with a NAD-dependent deacylase (9gmr) and a centromeric nucleosome complex (7k7g) (TM-score = 0.918), whereas the TM-score produced by US-align (0.414) would classify the pair as structurally unrelated. Similarly, GTcomplex accurately aligns assemblies containing nucleosomes bound to the transcription coactivator complex SAGA (6t9l) and the Dot1L methyltransferase (6nn6), achieving a TM-score of 0.764 compared with 0.385 for US-align, whose alignment misaligns part of the nucleosomes. A third example involving the DNA replication initiator protein DnaA (8btg) and a transcription factor pair (4xrs) illustrates the more accurate alignment produced by GTcomplex (0.541 vs. 0.463 by TM-score), revealing similar DNA-binding arrangements.

Examples from the NA-2-100 and NA-BA-2-100 datasets reveal improved chain mapping for nucleic acid complexes (Fig. 7d). For two synthetic DNA complexes (7u43 and 5i36), GTcomplex achieves a higher TM-score (0.830) than US-align (0.731). In another case, GTcomplex identifies a more complete correspondence between tRNAs interacting at different sites of the 70S ribosome by correctly mapping two tRNAs rather than one for the 1zo3-486d pair. Finally, GTcomplex produces a more accurate alignment between a prokaryotic DNA (7nbn) and a double-stranded RNA from bluetongue virus (1h1k), achieving a TM-score of 0.532 versus 0.440 for US-align and revealing structural similarities that may suggest similar allosteric signaling mechanisms.

### Case study highlights GTcomplex’s utility

A practical application of GTcomplex is illustrated through the analysis of an AlphaFold 3 model of a heterotrimer from Wenling crested flounder alphavirus (Model Archive [27] ID ma-6kpi5). The ma-6kpi5 model represents part of an asymmetric unit that would be used to construct the complete viral capsid, making the relative arrangement of its subunits critical for downstream architectural interpretation.

A GTcomplex search against PDB biological assemblies yielded more than 100 significant multimeric matches. Two of the top-scoring hits—representing the capsid of Getah virus (7fd2) and the asymmetric unit of Chikungunya virus (8fcg)—are shown in Fig. 7e. In both cases, the GTcomplex superpositions reveal that the terminal α-helices of the E1 and E2 glycoproteins in ma-6kpi5 adopt orientations that are inconsistent with those observed in homologous alphavirus structures. These deviations coincide with regions of low predicted confidence (pLDDT), suggesting that the helix orientations are likely incorrect.

This case study illustrates the use of GTcomplex to detect local modeling inaccuracies and guide structural interpretation in the context of experimentally determined macromolecular complexes.

## Discussion

GTcomplex provides a general, self-contained, and efficient solution for structure search and alignment of macromolecular complexes. Unlike commonly used approaches that rely on aligning individual chains followed by chain assignment, GTcomplex performs alignment at the level of whole complexes, with chain correspondence inferred directly from the global structural match. This design avoids heuristics tied to per-chain analyses and reduces errors that arise when the broader architectural context is ignored. By leveraging spatial indexing, GTcomplex achieves both high efficiency and high accuracy across diverse macromolecular architectures.

The benchmarking results underscore the limitations of methods that depend on pairwise chain comparisons or transformation-matrix clustering. In viral assemblies containing many similar or repeated chains, Foldseek-MM variants often misassign subunits and fail to recover global relationships, while the cost of numerous pairwise chain comparisons leads to substantial runtime inflation. GTcomplex, in contrast, maintains consistent accuracy and efficiency even for the largest assemblies.

To assess performance rigorously, we implemented a comprehensive evaluation framework based on cumulative metrics that measure alignment quality over all significant hits. This approach reveals subtle yet biologically meaningful differences between methods. Because TM-score values in the range [0.5; 1.0] correspond to significant fold-level similarity for proteins [28], extending this threshold to complexes after chain correspondence provides an interpretable indicator of sensitivity. Examining cumulative TM-score and RMSD trends further exposes tendencies toward overextended alignments: rapidly declining TM-scores accompanied by sharply increasing RMSDs indicate reduced accuracy and, consequently, lower sensitivity to structural similarity.

This region of divergence between methods, often emerging at a TM-score of approximately 0.5 (Figs. 2,4), is particularly informative for assessing sensitivity because it reflects the ability to detect structurally related complexes when similarities are limited. Consequently, aggregating performance across all significant hits and all queries captures a method’s ability to identify structural relationships in both similarity-rich and similarity-sparse settings, while reducing dependence on the composition of individual query sets. Complementing this analysis, the per-query sensitivity evaluation based on only a few top-ranked hits (Fig. 6) emphasizes queries with limited numbers of matches and independently confirms the conclusions drawn from the aggregate benchmarks.

The structural examples examined here further demonstrate that even modest improvements in alignment can affect biological interpretation. Because all subunits jointly define the architecture of a complex, they also constrain the search space for valid superpositions. Improvements in complex alignment therefore often stem from more accurate chain assignment. Correcting chain correspondences, especially in assemblies with high internal symmetry, may yield only moderate increases in TM-score, yet can produce more accurate and biologically relevant superpositions.

To support reproducible evaluation, we provide general scripts based on TM-align for computing similarity metrics for arbitrary complex alignments, enabling consistent and comparable assessment across methods.

Together, these results demonstrate that GTcomplex delivers state-of-the-art performance for complex alignment while remaining scalable and conceptually simple. Its general spatial-indexing framework applies naturally to proteins and nucleic acids and can be extended to complexes with small-molecule ligands, broadening the range of potential applications.

GTcomplex is distributed as a portable, user-friendly software package with support for large-scale clustering. It is also integrated into the GTalign webserver [29] (https://bioinformatics.lt/comer/gtalign), enabling high-throughput searches across the full PDB repository of biological assemblies. We anticipate that GTcomplex will facilitate evolutionary and functional studies, structural annotation, drug-discovery efforts, and advances in multimeric structure prediction [6, 30].

## Methods

### Algorithm specification

An overview of the GTcomplex algorithm is presented in the main text. A complete and formal specification of the algorithm is provided in Supplementary Section S2.

### GTcomplex software

GTcomplex is implemented using the CUDA architecture for GPU acceleration and is compatible with all modern NVIDIA GPU platforms. The software is distributed as open-source and includes precompiled binaries for Linux and Windows (x64), facilitating straightforward installation without requiring a CUDA toolchain. Source code and build instructions are provided to ensure full reproducibility and allow customization.

GTcomplex accepts input in PDB and PDBx/mmCIF formats, including gzip-compressed files, directories, and tar archives. Its command-line interface supports batch processing and deployment in high-performance, multi-GPU computing environments.

### GTcomplex web service

GTcomplex is integrated into the GTalign webserver [29], providing an interactive interface for structure search and alignment of both macromolecular monomers and complexes. The service currently supports searches against the complete PDB repository of biological assemblies.

### Structure representation

GTcomplex supports flexible specification of representative atoms for macromolecular complexes containing proteins and nucleic acids. By default, protein residues are represented by their Cα atoms and nucleotides by their C3✬ atoms. All experiments in this study were conducted using these default representations.

### Similarity prescreening

GTcomplex provides similarity prescreening controlled through the --pre-score option. When activated, candidate complex pairs with approximate similarity estimates (at --speed=16 after Stage 3) or with provisional TM-scores computed after Stage 4 (Fig. 1a) that fall below a user-defined threshold are excluded from further processing. Prescreening reduces runtime and operates independently of, and in addition to, the --speed parameter.

The default setting is --pre-score=0.4. For benchmarking, threshold values were adjusted to ensure adequate coverage across all evaluated --speed configurations: --pre-score=0.38 for the Ref-2-100, Full-PDB, Ref-BA-2-100, and ProtNA-BA-100 datasets, --pre-score=0.3 for Viral-C, --pre-score=0.2 for NA-2-100, and --pre-score=0.15 for NA-BA-2-100. Benchmarking results for different prescreening thresholds are provided in Supplementary Figs. S6–S7 (Viral-C) and Supplementary Figs. S8–S9 (NA-2-100).

### Alignment accuracy evaluation

Alignment accuracy is assessed by evaluating how well two macromolecular complexes match in 3D space after one complex is superimposed onto the other according to the alignment being evaluated.

We employed the TM-align [24] and TM-score [21] programs to calculate TM-score, RMSD, and GDT TS that provide standardized measures of the spatial agreement between structures under a specified alignment. TM-align provides the -I option, which restricts TM-score optimization to a user-provided alignment, ensuring that the resulting optimal superposition is constrained by that alignment. However, these tools operate only on single-chain structures.

To evaluate complex-level alignments, the structure files of each complex are first reordered according to the chain assignment produced by the method under evaluation. The reordered complexes, together with the concatenated chain-to-chain alignments, are then provided to TM-align or TM-score as if each complex were a single-chain structure (Fig. 8). This procedure provides a uniform framework for evaluating alignment accuracy.

**Fig. 8.**
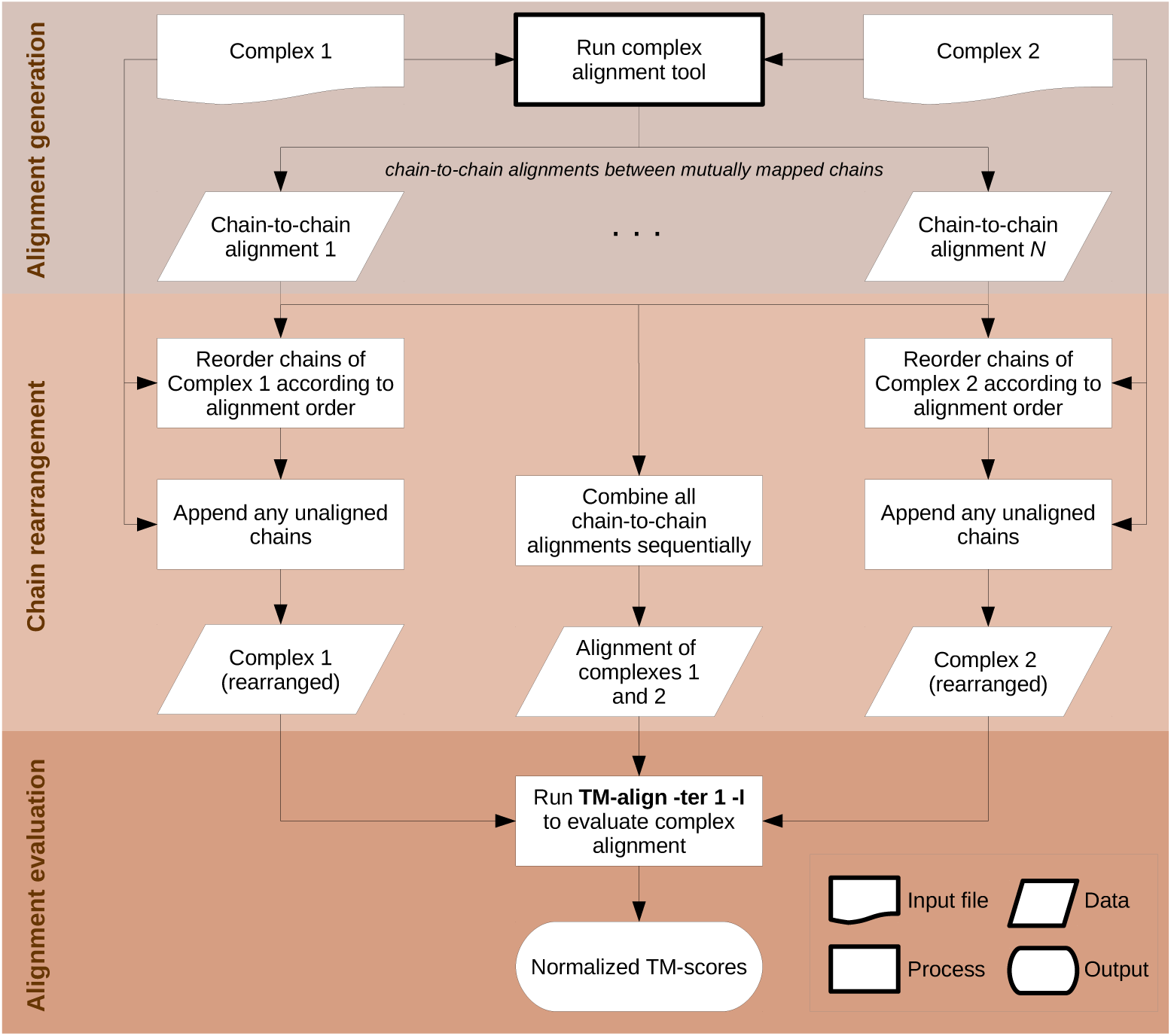
Flowchart of the procedure for evaluating alignment accuracy for macromolecular complexes. The diagram illustrates how TM-scores and RMSD are calculated using TM-align for a complex alignment represented as multiple chain-to-chain alignments. The same procedure is used to calculate GDT TS, substituting the adapted TM-score program for TM-align.

A standardized evaluation pipeline is essential for ensuring comparability across tools and for avoiding differences introduced by method-specific scoring implementations. For example, Foldseek-MM variants compute approximate TM-scores for speed, making their reported values not directly comparable to scores from methods such as US-align. However, Supplementary Fig. S10 shows that the results remain stable when each method’s own TM-scores are used to rank their alignments.

We introduced minimal modifications to the TM-score source code to compute normalized GDT TS scores constrained by a given alignment, and also we introduced two minor changes (commented lines) to the TM-align code (TM-align RNA) to support TM-score calculations for nucleic acid structures [31]. Both adapted tools are publicly available.

Finally, we verified that the statistics computed by this unified procedure match those produced directly by US-align. TM-scores and RMSDs reported by US-align were used as-is in all experiments and not recalculated. Scores were also used as reported for protein-nucleic acid assembly alignments (ProtNA-BA-100).

### PDB clustering for dataset construction

The entire PDB was clustered to construct the Ref-2-100 dataset of structurally diverse protein complexes using GTcomplex at a TM-score threshold of 0.5 and a minimum coverage of 70%. Clustering was performed on NVIDIA Saturn Cloud with the command:

gtcomplex -v --cls=mmCIF -o <output_dir> --speed=9 --pre-score=0.4 --cpu-threads-reading =20 --dev-N=8 --cls-algorithm=0 --cls-coverage=0.7 --cls-threshold=0.5 -c <cache_dir>

where mmCIF represents the directory containing PDB structures in PDBx/mmCIF format downloaded on 27 June 2025. The resulting cluster list is publicly available.

This clustering procedure took five days on eight NVIDIA A100 GPUs (--dev-N=8) using GTcomplex version 1.0.0. Subsequent releases, including version 1.0.9 used for all benchmarking in this study, introduced substantial performance improvements, yielding nearly an order-of-magnitude reduction in runtime. As a result, clustering with version 1.0.9 or later is expected to be significantly faster.

The query and subject structure files derived from the resulting clusters were uncompressed, and only their first models were used in the experiments.

For the ProtNA-BA-100 dataset, protein-nucleic acid assemblies were clustered using GTcomplex with parameters --speed=12 --pre-score=0.4 --cls-coverage=0.7 and --cls-threshold=0.5. Clustering was performed on a single NVIDIA GeForce RTX 4090 Laptop GPU and completed in 6 h. The resulting cluster list is publicly available.

### System configuration for benchmarking

Benchmark tests on the Ref-2-100, Full-PDB, Viral-C, and NA-2-100 datasets were performed on NVIDIA Saturn Cloud. Benchmarks were executed on an instance with an AMD EPYC 7J13 64-core CPU (3.2 GHz boost clock), 1 TB RAM, and four NVIDIA A100-SXM4-80GB GPUs.

Benchmark tests on the Ref-BA-2-100, ProtNA-BA-100, and NA-BA-2-100 datasets were performed on the Vast.ai cloud platform using an AMD EPYC 7J13 64-core CPU (configured by the resource provider at a 2.45 GHz base clock), 132 GB RAM, and two NVIDIA A100-SXM4-40GB GPUs.

All CPU-based tools were run using all 64 CPU cores. GTcomplex was configured (--dev-N=2) to use two A100 GPUs to provide a balanced comparison relative to CPU-based methods.

Table 1 reports GTcomplex runtimes across all evaluated --speed configurations for each dataset, obtained using two A100 GPUs and, for comparison, a single desktop-grade NVIDIA GeForce RTX 5090 (32 GB). The results show that a single modern desktop GPU delivers performance comparable to two older server-grade GPUs.

**Table 1.** Runtimes (in seconds) for GTcomplex across different --speed configurations and hardware platforms for each dataset

|  | 2×A100<br>(*Server*) | 1×GeForce RTX 5090<br>(*Desktop*) |
| --- | --- | --- |
| *Ref-2-100* |  |  |
| GTcomplex --speed=0 | 3801.4 | 3920.7 |
| GTcomplex --speed=4 | 3456.5 | 3650.3 |
| GTcomplex --speed=9 | 2149.2 | 2200.1 |
| GTcomplex --speed=10 | 1376.7 | 1442.2 |
| GTcomplex --speed=12 | 615.4 | 594.3 |
| GTcomplex --speed=16 | 195.0 | 205.4 |
| *Full-PDB* |  |  |
| GTcomplex --speed=0 | 18036.4 | 18340.6 |
| GTcomplex --speed=4 | 14948.5 | 15719.2 |
| GTcomplex --speed=9 | 10622.4 | 11014.5 |
| GTcomplex --speed=10 | 6612.5 | 7037.6 |
| GTcomplex --speed=12 | 3399.3 | 3369.1 |
| GTcomplex --speed=16 | 1088.0 | 1069.5 |
| *Viral-C* |  |  |
| GTcomplex --speed=0 | 532.0 | 438.3 |
| GTcomplex --speed=4 | 263.1 | 234.2 |
| GTcomplex --speed=9 | 100.6 | 94.2 |
| GTcomplex --speed=10 | 81.3 | 78.6 |
| GTcomplex --speed=12 | 38.9 | 31.5 |
| GTcomplex --speed=16 | 25.3 | 19.5 |
| *NA-2-100* |  |  |
| GTcomplex --speed=0 | 50.9 | 52.0 |
| GTcomplex --speed=4 | 39.8 | 39.8 |
| GTcomplex --speed=9 | 43.6 | 43.6 |
| GTcomplex --speed=10 | 28.6 | 29.5 |
| GTcomplex --speed=12 | 19.8 | 19.4 |
| GTcomplex --speed=16 | 11.4 | 12.7 |

GPU memory capacity above 16 GB does not affect GTcomplex runtime performance. Combined with the--dev-mem option, which specifies the amount of GPU memory allocated per process, this enables running multiple GTcomplex processes concurrently on a single GPU. For example, on an A100-SXM4-80GB GPU, setting --dev-mem=16384 (16 GB) permits up to five concurrent GTcomplex processes.

### Runtime evaluation

Runtimes for all tools were measured using the Linux time command.

### GTcomplex command-line configuration

All analyses were performed using GTcomplex version 1.0.9. The general command used to perform GTcomplex searches was

<p>gtcomplex -v --qrs=<query_dir> --rfs=<db_dir> -o <output_dir> -c <cache_dir> --pre-score =<prescore_threshold> --speed=<speed> --dev-N=2 -s <output_threshold> --nhits=<n_hits> --nalns=<n_hits></p>

where <query_dir> and <db_dir> denote the directories containing the query and subject structures, respectively. For the Viral-C dataset, both directories were identical. <output_dir> specifies the output directory, and <cache_dir> specifies a temporary directory used to store cached data when repeated file reads become more expensive than computation.

<prescore_threshold> denotes the similarity-prescreening cutoff. For the main benchmarking results, it was set to 0.38 for the Ref-2-100, Full-PDB, Ref-BA-2-100, and ProtNA-BA-100 datasets, 0.3 for Viral-C, 0.2 for NA-2-100, and 0.15 for NA-BA-2-100. (Supplementary Figs. S6–S9 report results across three thresholds for Viral-C and NA-2-100.) <speed> denotes the evaluated --speed configurations. <output_threshold> specifies the minimum TM-score for reporting alignments and was set to 0.4 and 0.3 (--speed={12,16}) for Ref-2-100, Full-PDB, Ref-BA-2-100, and ProtNA-BA-100, to 0 for Viral-C, and to 0.2 for NA-2-100 and NA-BA-2-100. <n_hits> controls the maximum number of alignments returned per query and was set to 4000 for Ref-2-100 and Ref-BA-2-100, 20000 and 10000 (--speed={12,16}) for Full-PDB, 8000 for ProtNA-BA-100, and left at the default value of 2000 for Viral-C, NA-2-100, and NA-BA-2-100.

Additional options were used for Viral-C, NA-2-100, and NA-BA-2-100. For Viral-C,--dev-queries-max-chains=420 and --dev-queries-total-length-per-chunk=42000 enabled processing of capsids with up to 420 subunits and total residue lengths up to 42,000. For NA-2-100 and NA-BA-2-100, --cpu-threads-reading=2 was used to reduce the number of file-reading threads, thereby increasing computational batch size.

### US-align configuration

US-align version 20241108 was used for all benchmark comparisons. The tool was executed using the following command:

USalign <query_file> <subject_file> -mm 1 -ter 1 [-fast]

where <query_file> and <subject_file> denote the input structures. The -fast option was used to run US-align in its fast mode.

Searches for each dataset were parallelized across 64 CPU cores by distributing query complexes over available workers. For the Full-PDB, Ref-BA-2-100, and ProtNA-BA-100 datasets, the fast mode was omitted due to long runtimes. For Full-PDB, the standard mode was executed with the alignment-free output format (-outfmt 2) to limit disk usage, which consequently made GDT TS evaluation for this dataset unavailable.

### Foldseek-MM configuration

Foldseek version 1085e5b0f282b57a91ad3920074ab7a652987ff5 (downloaded on 22 July 2025) was used for benchmarking. Searches were performed using the command:

foldseek easy-multimersearch <query_dir> <db_dir> <output_file> <tmp_dir> --threads 64--max-seqs <n_hits> --format-output query,target,evalue,bits,qtmscore,ttmscore, alntmscore,rmsd,complexqtmscore,complexttmscore,complexu,complext,complexassignid, qstart,qend,qlen,tstart,tend,tlen,qaln,taln

where <query_dir> and <db_dir> specify the directories containing the query and subject structures, respectively. <output_file> denotes the output file, and <tmp_dir> a temporary working directory.<n_hits> controls the maximum number of complex-level alignments reported per query and was set to 4000 for Ref-2-100 and Ref-BA-2-100, 8000 for Full-PDB, and left at the default value for Viral-C.

Foldseek-MM-TM results were obtained by enabling TM-align-based chain-level alignment with the--alignment-type 1 option.

Prior to benchmarking, dry runs were performed to preload large datasets into memory and reduce I/O overhead during searches.

Although Foldseek supports GPU acceleration for the sequence prefiltering stage, enabling GPU mode for complex alignment resulted in only modest speed gains while introducing pervasive inaccuracies in complex alignments. Consequently, GPU acceleration was not used for Foldseek-based complex alignment.

### Figure preparation

Molecular graphics were generated using UCSF ChimeraX version 1.3 [32]. Plots were produced using the ggplot2 package [33] in R [34] (versions 3.6.0 and 4.4.2).

## Data availability

All datasets were derived from the Protein Data Bank (PDB). Structure files are available from the RCSB PDB at https://www.rcsb.org/downloads. The benchmark data generated in this study are available at https://github.com/minmarg/gtcomplex-evaluation.

## Code availability

GTcomplex source code and software packages are available at https://github.com/minmarg/gtcomplex. Corresponding releases are archived on Zenodo at https://zenodo.org/records/17937811 (version 1.0.9; doi: 10.5281/zenodo.17937811) and https://zenodo.org/records/17937810 (versions >1.0.9; doi: 10.5281/zenodo.17937810). Lists of structure identifiers together with programs and scripts used for benchmarking and figure generation (plots) are available at https://github.com/minmarg/gtcomplex-evaluation.

## Acknowledgements

This work was supported by an NVIDIA Academic Grant (M.M.) and by Research Council of Lithuania (LMTLT) grant S-MIP-23-104 (M.M.). The project utilized 8x NVIDIA A100 GPUs.

The author thanks Justas Dapkūnas for adapting the GTalign webserver [29] frontend to support GTcomplex and for suggesting the PDB biological assemblies database.

## Author contributions

M.M. conceived and supervised the project, designed the algorithms, developed GTcomplex, performed the analyses, and wrote the manuscript.

## Competing interests

M.M. is a co-founder of Hiomics digital and declares non-financial competing interests.

## S1 Supplementary results

### S1.1 Supplementary figures

**Fig. S1.**
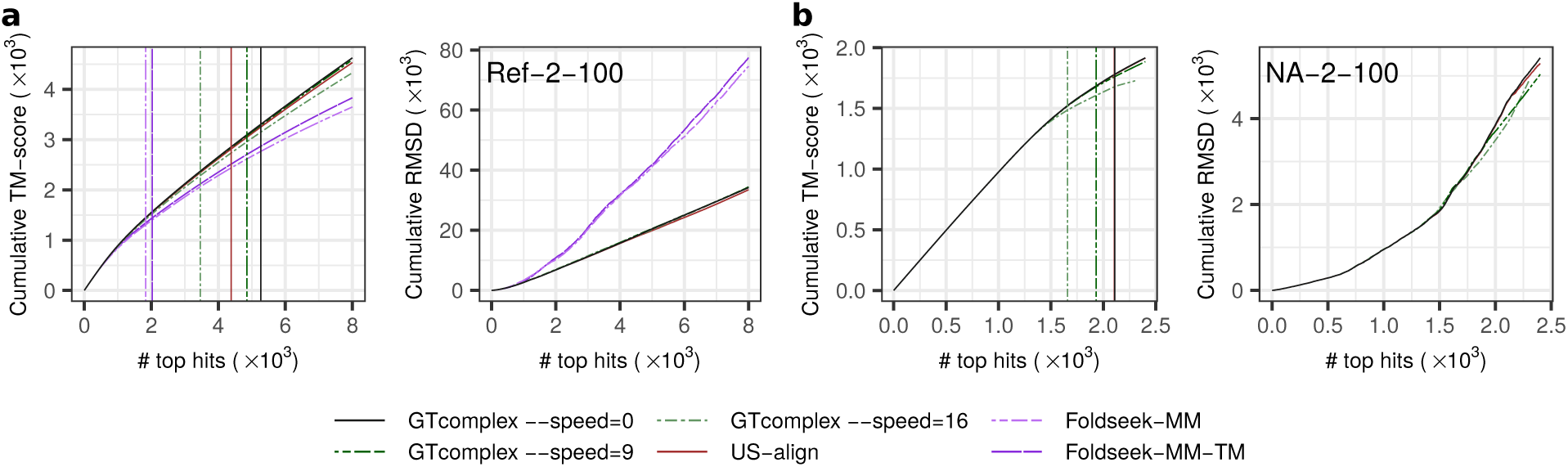
Benchmarking results on the Ref-2-100 (a) and NA-2-100 (b) datasets using the higher-scoring of two reciprocal alignments for each query-subject pair. Left and right panels show cumulative TM-score and RMSD, respectively, across top-ranked alignments. Vertical lines indicate the number of alignments with TM-score ≥ 0.5. Alignments are ranked by TM-score normalized by the length of the shorter complex. TM-scores correspond to TM-align recalculations of the alignments.

**Fig. S2.**
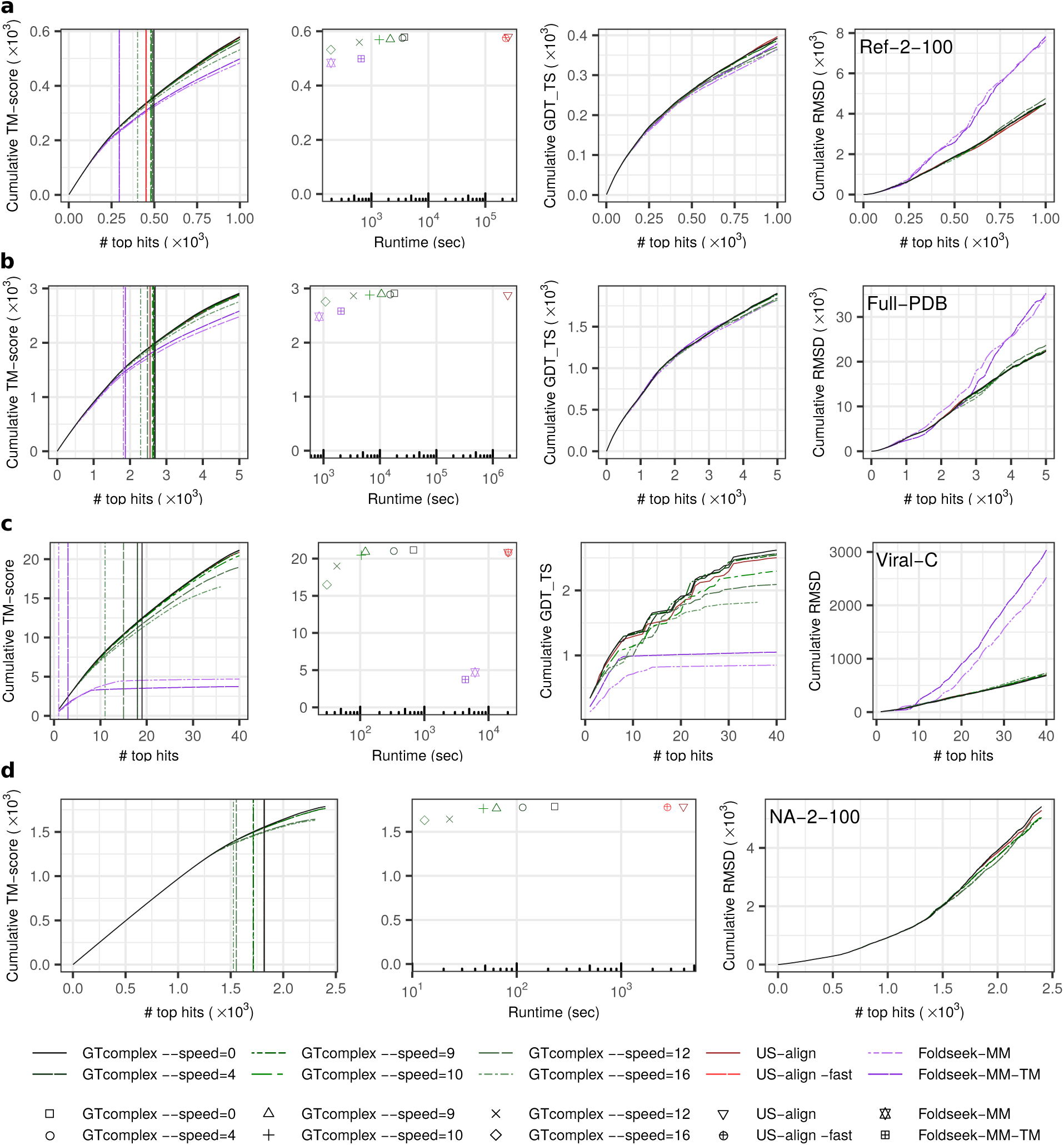
Benchmarking results on the Ref-2-100 (a), Full-PDB (b), Viral-C (c), and NA-2-100 (d) datasets, evaluated using TM-score and GDT_TS metrics normalized by query length. Left panels show cumulative TM-score as a function of the number of top-ranked alignments (ranked by TM-score). Vertical lines indicate the number of alignments with TM-score ≥ 0.5. Middle-left panels in (a–c) and the central panel in (d) plot cumulative TM-score versus runtime (seconds). Middle-right panels in (a–c) plot cumulative GDT_TS, and right panels plot cumulative RMSD, each as a function of the number of top-ranked alignments. Alignments are ranked by TM-score normalized by the length of the query complex. In the left, middle-left (a–c), and central (d) panels, the TM-scores correspond to TM-align recalculations of the alignments, and the ranking of alignments is based on these recalculated TM-scores.

**Fig. S3.**
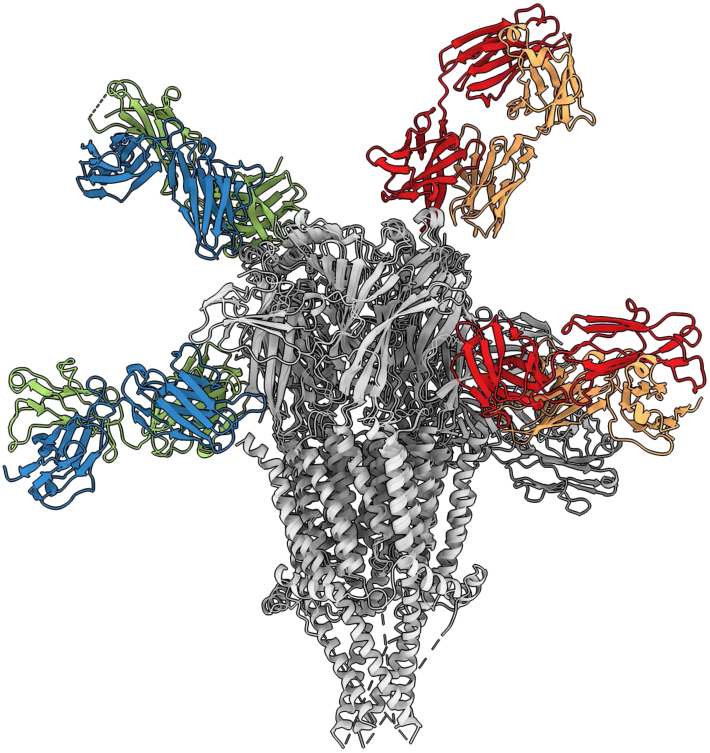
Example of spatial mismatches produced by Foldseek-MM and Foldseek-MM-TM. The figure shows the superimposition of complexes 8ssz and 9gu1 as aligned by Foldseek-MM-TM. Spatially distant chains that are incorrectly paired and aligned, resulting in a large RMSD of 36.60 Å (36.96 Å for Foldseek-MM), are highlighted in matching colors. All other chains are shown in gray.

**Fig. S4.**
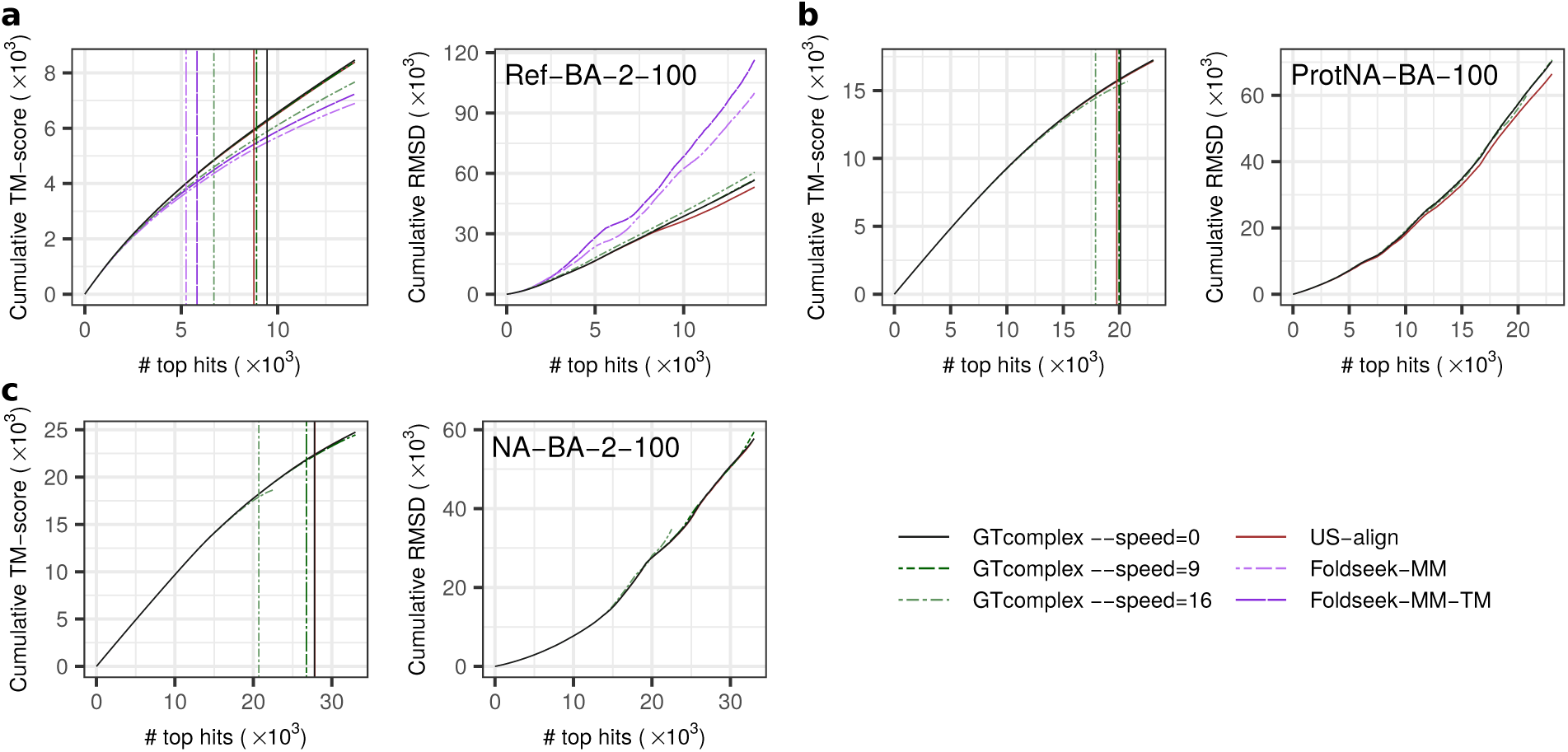
Benchmarking results on the Ref-BA-2-100 (a), ProtNA-BA-100 (b), and NA-BA-2-100 (c) datasets using the higher-scoring of two reciprocal alignments for each query-subject pair. Left and right panels show cumulative TM-score and RMSD, respectively, across top-ranked alignments. Vertical lines indicate the number of alignments with TM-score ≥ 0.5. Alignments are ranked by TM-score normalized by the length of the shorter assembly. TM-scores are recalculated using TM-align as described in Methods, except for ProtNA-BA-100 (b).

**Fig. S5.**
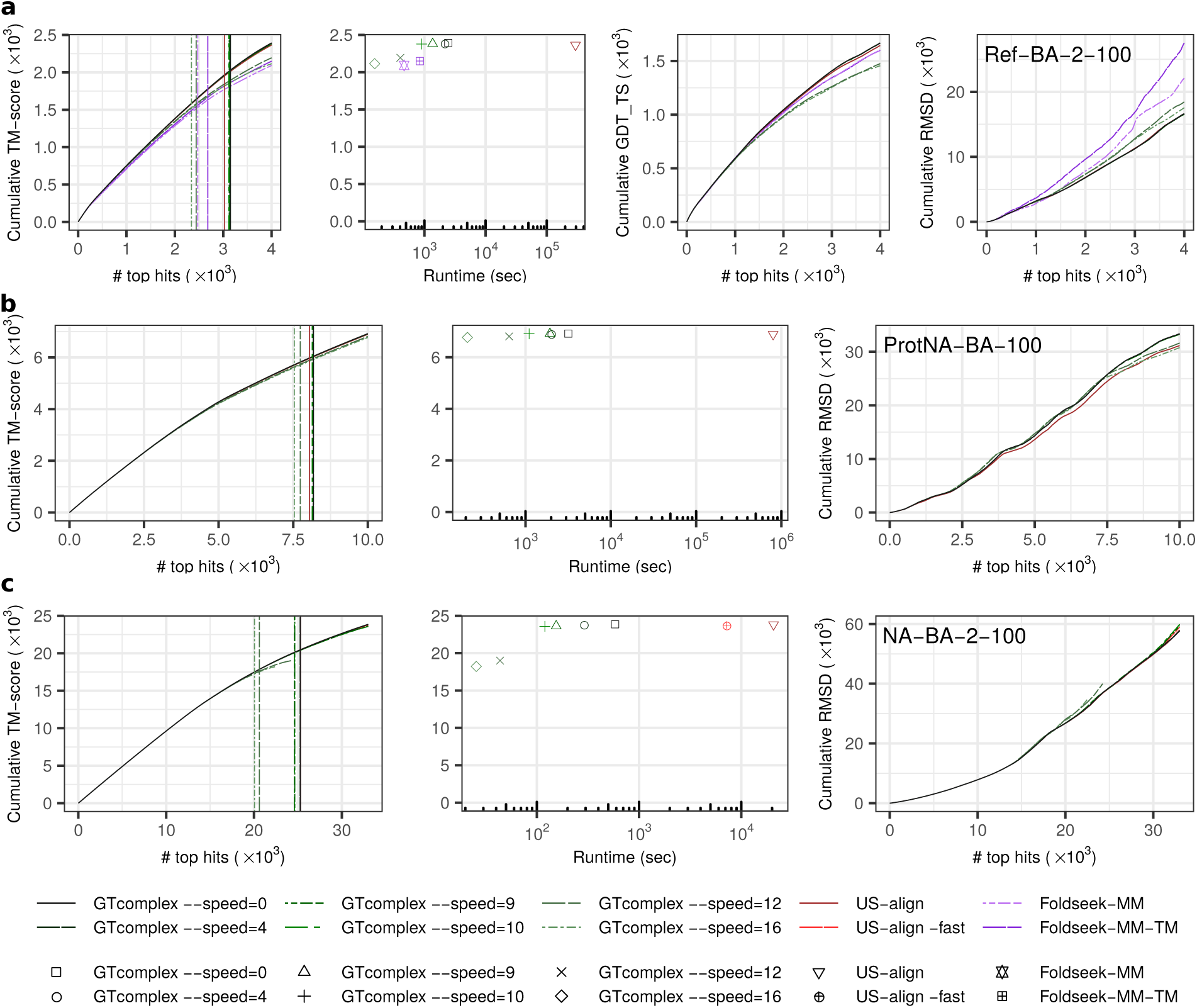
Benchmarking results on the Ref-BA-2-100 (a), ProtNA-BA-100 (b), and NA-BA-2-100 (c) biological assembly datasets using TM-score and GDT TS normalized by query length. Left panels show cumulative TM-score across top-ranked alignments. Vertical lines indicate the number of alignments with TM-score ≥ 0.5. The middle-left panel in (a) and the central panels in (b,c) show cumulative TM-score as a function of runtime. The middle-right panel in (a) shows cumulative GDT TS, and the right panels show cumulative RMSD across top-ranked alignments. TM-scores are recalculated using TM-align as described in Methods, except for ProtNA-BA-100 (b).

**Fig. S6.**
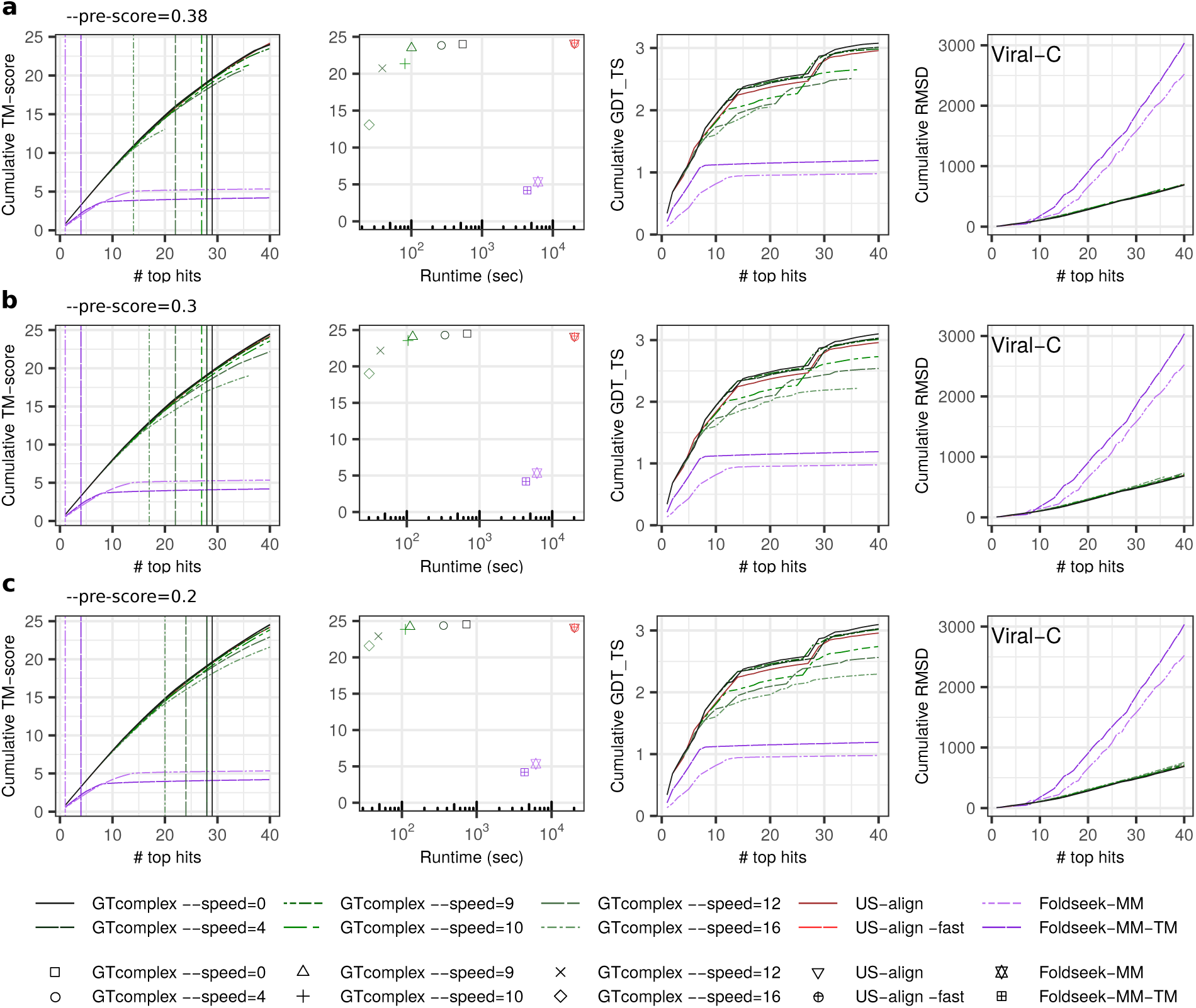
Benchmarking results on the Viral-C dataset for three different prefiltering settings: --pre-score=0.38 (a),--pre-score=0.3 (b), and --pre-score=0.2 (c). Alignments are ranked by TM-score normalized by the length of the shorter complex. Panel definitions follow those in Fig. 2 of the main text.

**Fig. S7.**
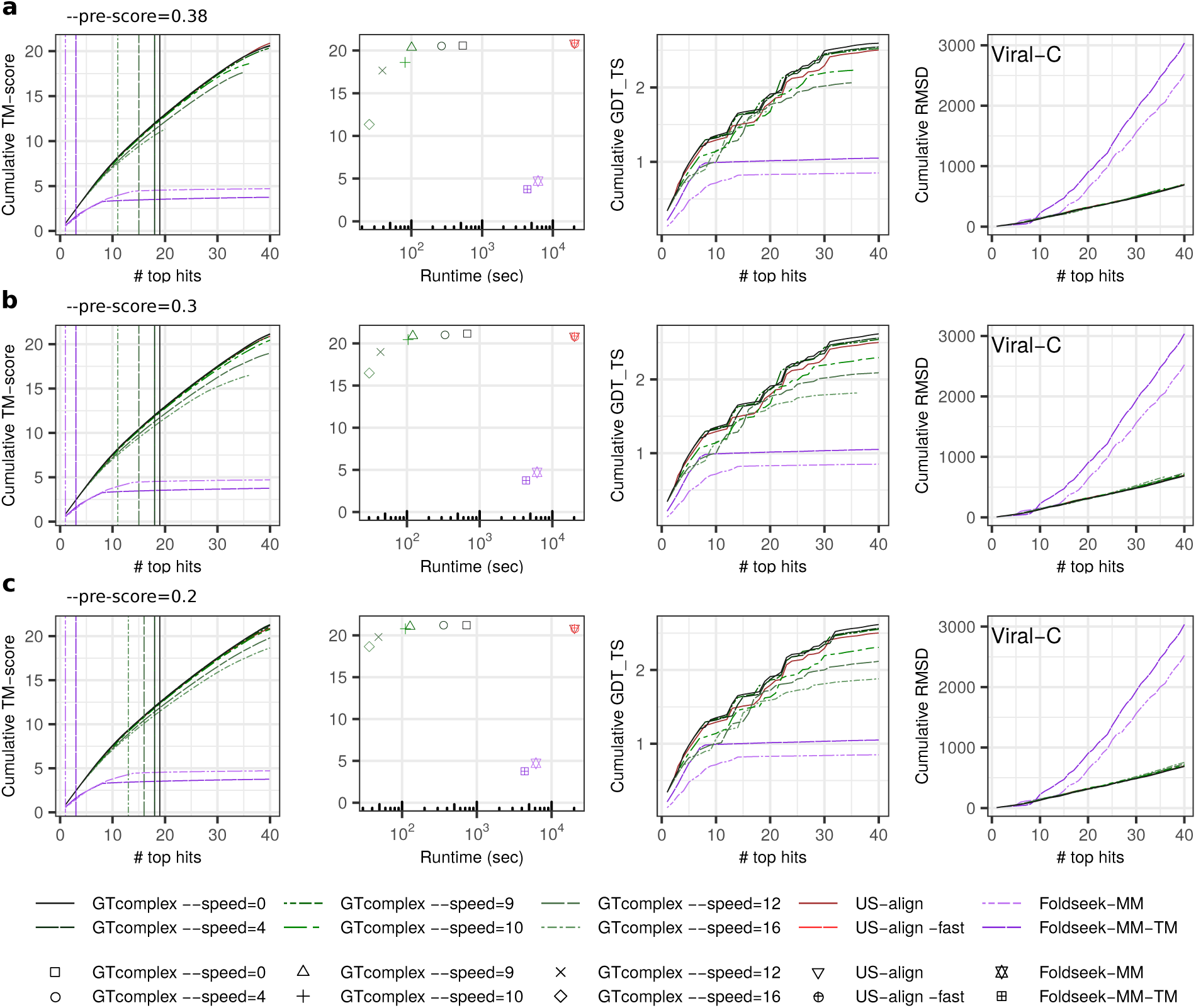
Benchmarking results on the Viral-C dataset for three different prefiltering settings: --pre-score=0.38 (a),--pre-score=0.3 (b), and --pre-score=0.2 (c). Alignments are ranked by TM-score normalized by the length of the query complex. Panel definitions follow those in Fig. S2.

**Fig. S8.**
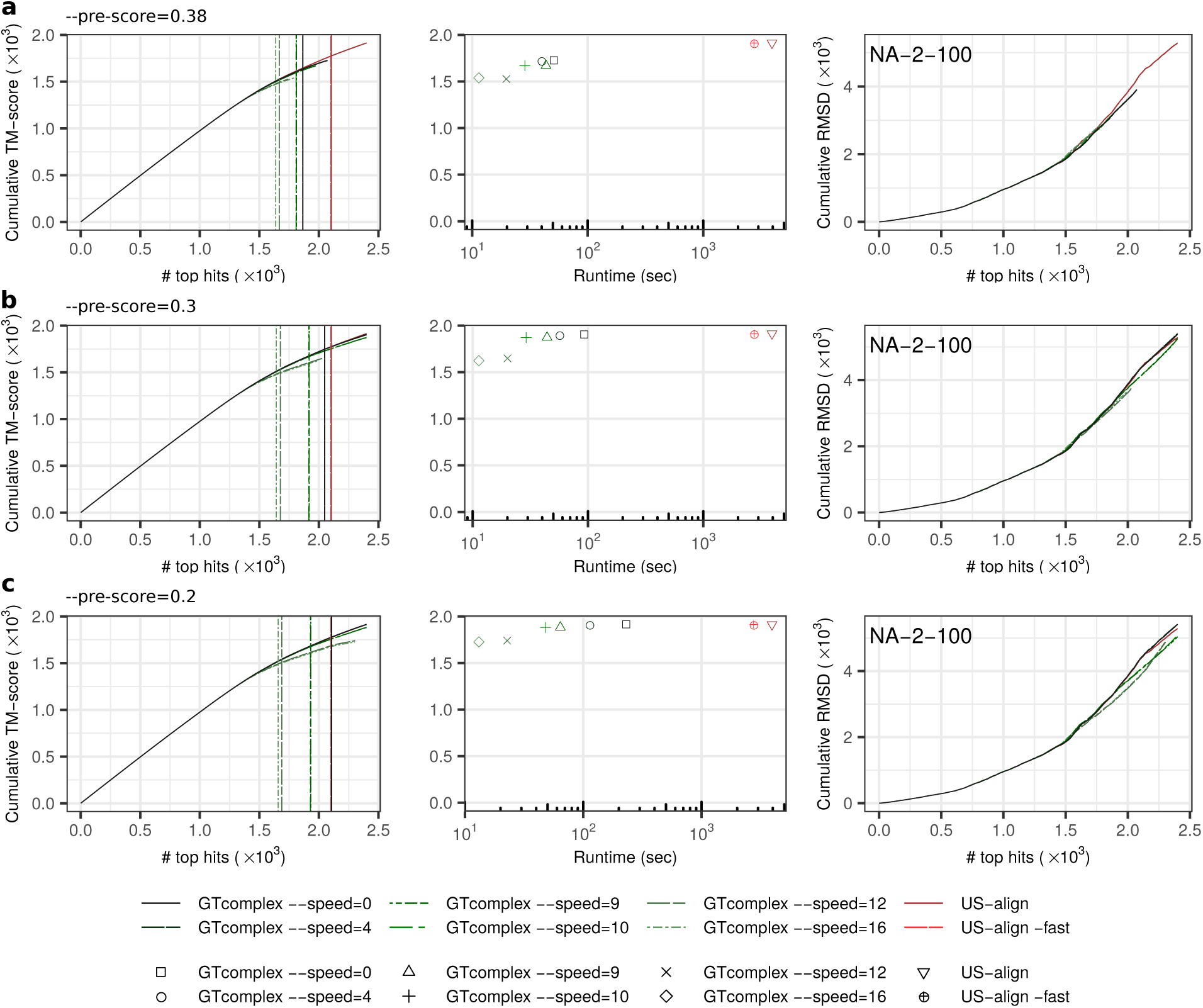
Benchmarking results on the NA-2-100 dataset for three different prefiltering settings: --pre-score=0.38 (a), --pre-score=0.3 (b), and --pre-score=0.2 (c). Alignments are ranked by TM-score normalized by the length of the shorter complex. Panel definitions follow those in Fig. 2d of the main text.

**Fig. S9.**
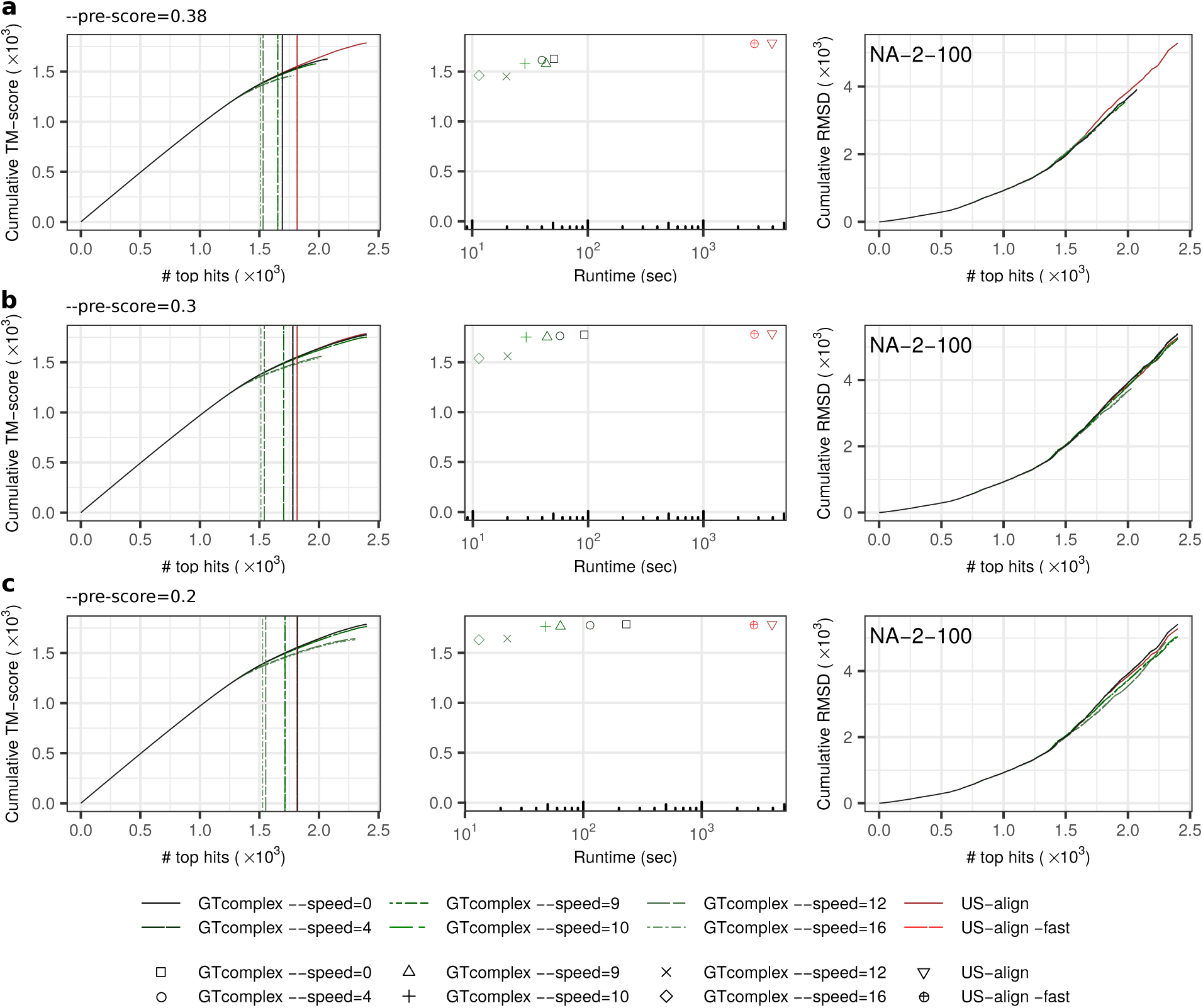
Benchmarking results on the NA-2-100 dataset for three different prefiltering settings: --pre-score=0.38 (a),--pre-score=0.3 (b), and --pre-score=0.2 (c). Alignments are ranked by TM-score normalized by the length of the query complex. Panel definitions follow those in Fig. S2d.

**Fig. S10.**
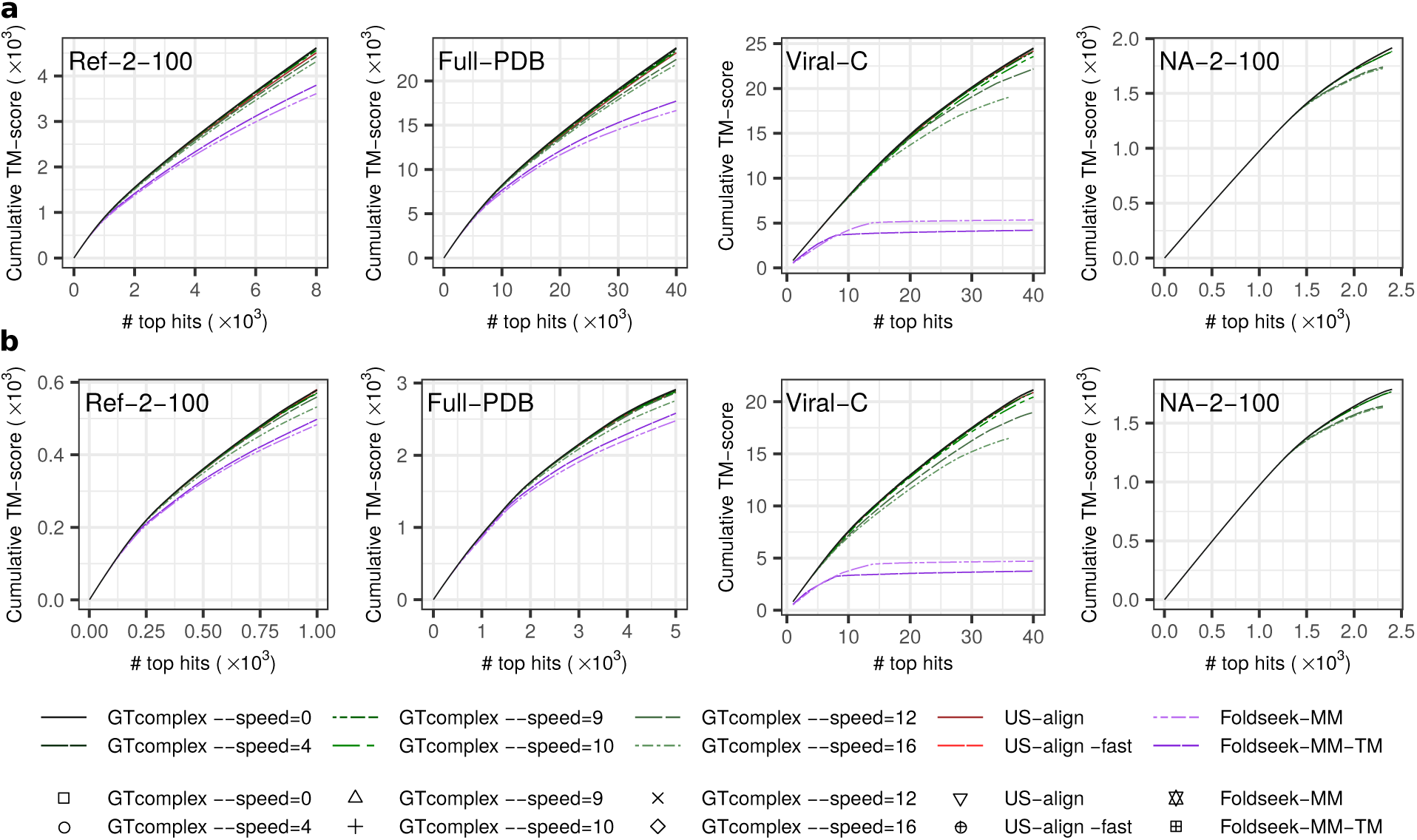
Rate of accurate alignments, evaluated by cumulative TM-score as a function of the number of top-ranked alignments for the Ref-2-100, Full-PDB, Viral-C, and NA-2-100 datasets. Unlike the left panels of Fig. 2 in the main text and Fig. S2, here TM-scores computed by each tool are used to rank alignments. a, Alignments ranked by TM-score normalized by the length of the shorter complex. b, Alignments ranked by TM-score normalized by the length of the query complex.

## S2 Supplementary methods

### S2.1 Efficient identification of optimal superpositions

This section describes the core algorithms that enable GTcomplex to identify optimal structural superpositions of macromolecular complexes. To efficiently explore vast superposition space and converge on the most consistent superpositions, GTcomplex employs spatial indexing to generate candidate transformations, followed by iterative alignment, chain assignment, and refinement steps. Highly customizable, these algorithms enable sensitive and scalable structure alignment across large datasets.

The top-level algorithm is presented in Algorithm 1. It accepts as input two batches of macromolecular complex structures, associated structural descriptors, and user-defined parameters, and it produces as output optimal superpositions, expressed as rigid-body transformation matrices (R, t), for all pairs of structures drawn from distinct batches. The two batches, representing the sets of query and subject structures, are processed in parallel.

In the algorithmic specification, bold lowercase letters denote vectors, and italic uppercase letters denote multi-dimensional matrices or collections. The operator [·]_mut_ indicates a mutually exclusive operation preventing concurrent access to the same data by different threads.

**Algorithm 1.**
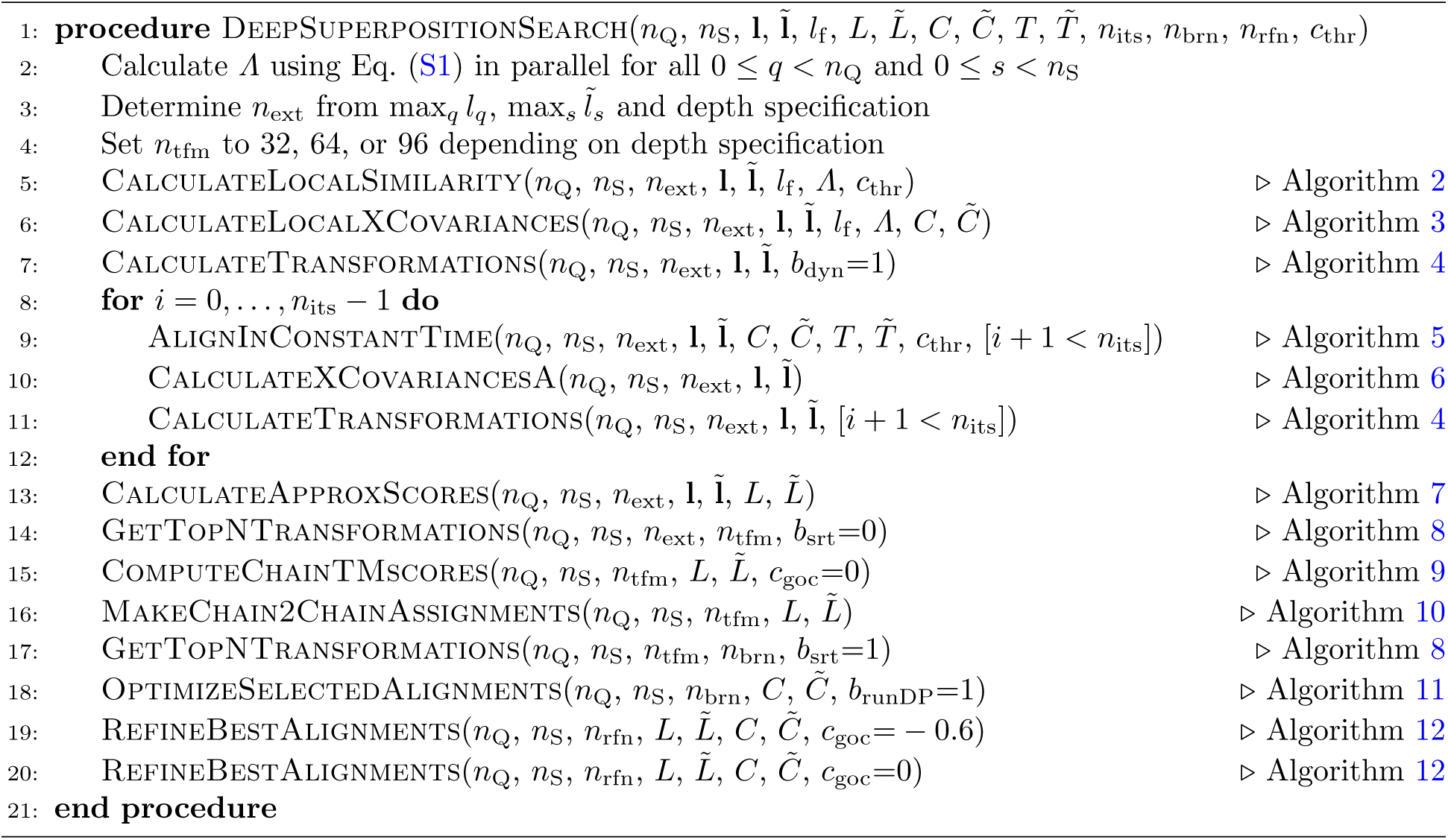
Find optimal structural superpositions through deep search.

The parameters used in Algorithm 1 are defined as follows. n_Q_ and n_S_ denote the numbers of query and subject complex structures in their respective batches. l and ̃l represent the lengths of the query and subject complexes, respectively. l_f_ specifies the context size used for local similarity evaluation and the generation of initial local alignment-based superpositions.

L and ̃L denote the chain-level lengths of the query and subject complexes, respectively, where L*_q, hq_* is the length of chain h*_q_* of query complex q, and ̃L*_̃s,h_* is defined analogously for subject complex s. C and ̃C represent the Cartesian coordinates of the query and subject structures. Specifically, C*_q,pq_* ∈ R^3×1^ denotes the coordinates of residue or nucleotide p*_q_* in query structure q, and ̃C*_s,p,̃s_* ∈ R^3×1^ has the corresponding meaning for subject structure s. The position indices p*_q_* and ̃p;*_s_* increase continuously along the entire complexes, spanning all constituent chains.

T and ̃T represent secondary structure assignments for protein query and subject complexes or nucleotide sequences for nucleic acid complexes. The complex type, protein or nucleic acid, is determined based on the predominant atom type, i.e., whether amino acids or nucleotides constitute the majority of the structure.

n_its_, n_brn_, and n_rfn_ are configurable parameters representing the number of deep, spatial index-driven superposition search iterations, top-performing superposition branches explored in detail, and refinement rounds, respectively. The parameter c_thr_ defines the local structural similarity threshold that triggers superposition analysis.

The algorithm operates on complete macromolecular complexes rather than treating them as separate chains. Chain-to-chain assignments are subsequently derived from the optimal complex-level superpositions identified during the search.

The algorithm begins by computing a dynamic programming (DP) matrix of local scores (line 2) using the following recursive relation:

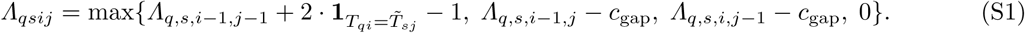

The one-byte variables Λ*_qsij_*are capped at a maximum value of 252, with the two least significant bits reserved for backtracking information. Initial superpositions (lines 6–7) are then identified based on structural regions exhibiting high local similarity (line 5). The total number of regions, n_ext_, is determined by the lengths of the structures, and the regions are distributed evenly along them.

After the initial superpositions are computed, several rounds of constant-time, spatial index-driven alignment are performed to generate the corresponding superpositions (lines 9–11). These steps form the core of the algorithm, treating each position independently and enabling rapid exploration of the superposition space across different configurations.

The atom-independent structural analysis produces sequence-order-independent alignments that capture spatial correspondence but not topological similarity. To account for structural topology, a post-processing step is applied.

The post-processing step (line 13) converts sequence-order-independent alignments into approximate order-dependent TM-scores computed in sublinear time. A small number of transformation matrices (n_tfm_) with the highest approximate scores are then selected (line 14) to calculate chain-level TM-scores (line 15) using the parallel COMER2 DP algorithm [1]. Note that only complex-level transformation matrices are used throughout all algorithmic stages.

After determining chain-to-chain correspondences by maximizing the sum of chain-level TM-scores (line 16), a smaller number of transformation matrices (n_brn_ < n_tfm_) corresponding to the highest TM-scores are selected (line 17) for further optimization of the associated structural alignments (line 18). Finally, the best-performing alignments—each representing a unique query-subject pair—are refined through iterative complex alignment, chain reassignment, and optimization of the resulting complex alignments (lines 19–20).

Algorithm 1 produces transformation matrices for each query-subject complex pair, which are subsequently used in the final stages for fine-grained alignment refinement. The procedure for such refinement is summarized in Algorithm 12.

**Algorithm 2.**
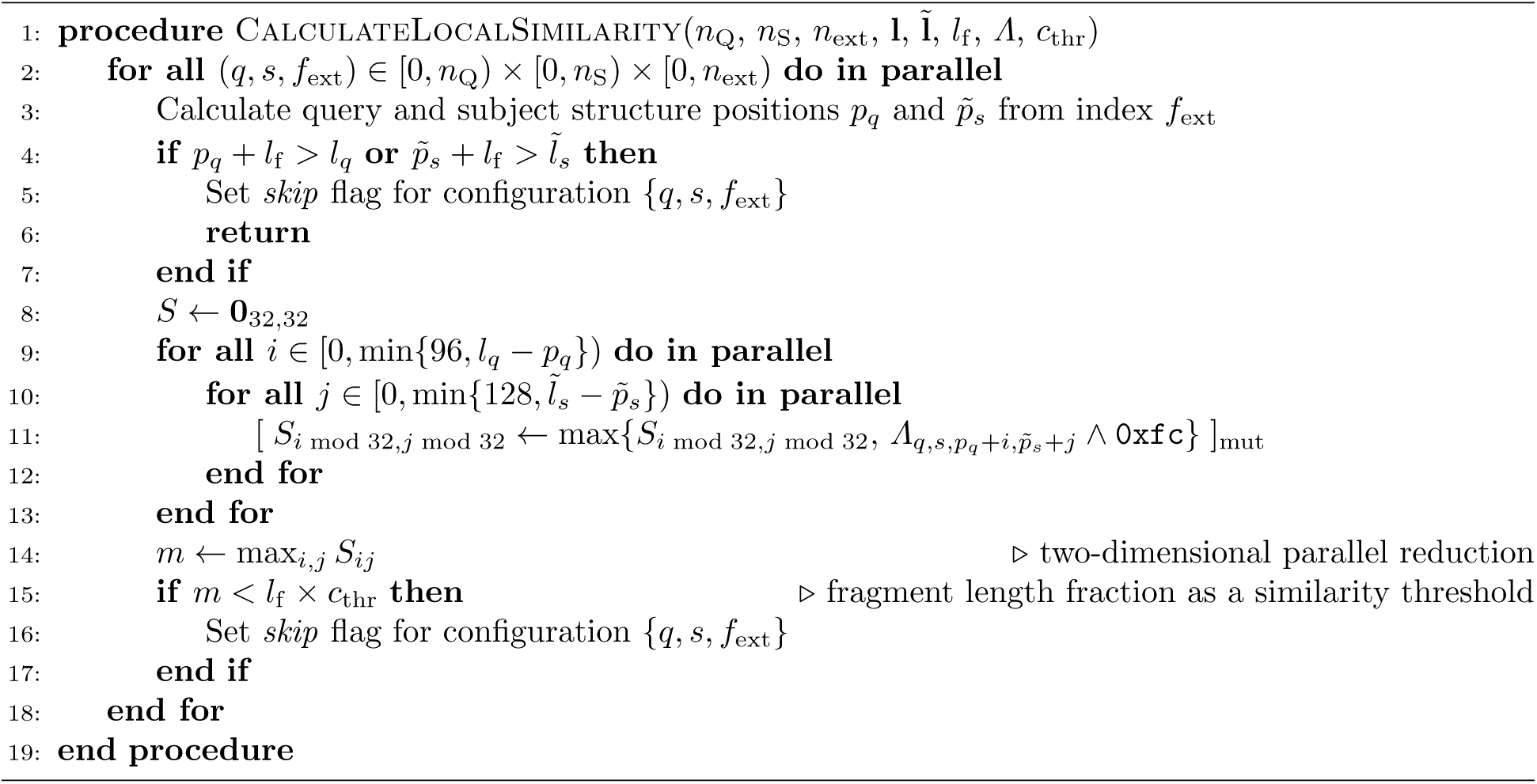
Calculate local similarity.

**Algorithm 3.**
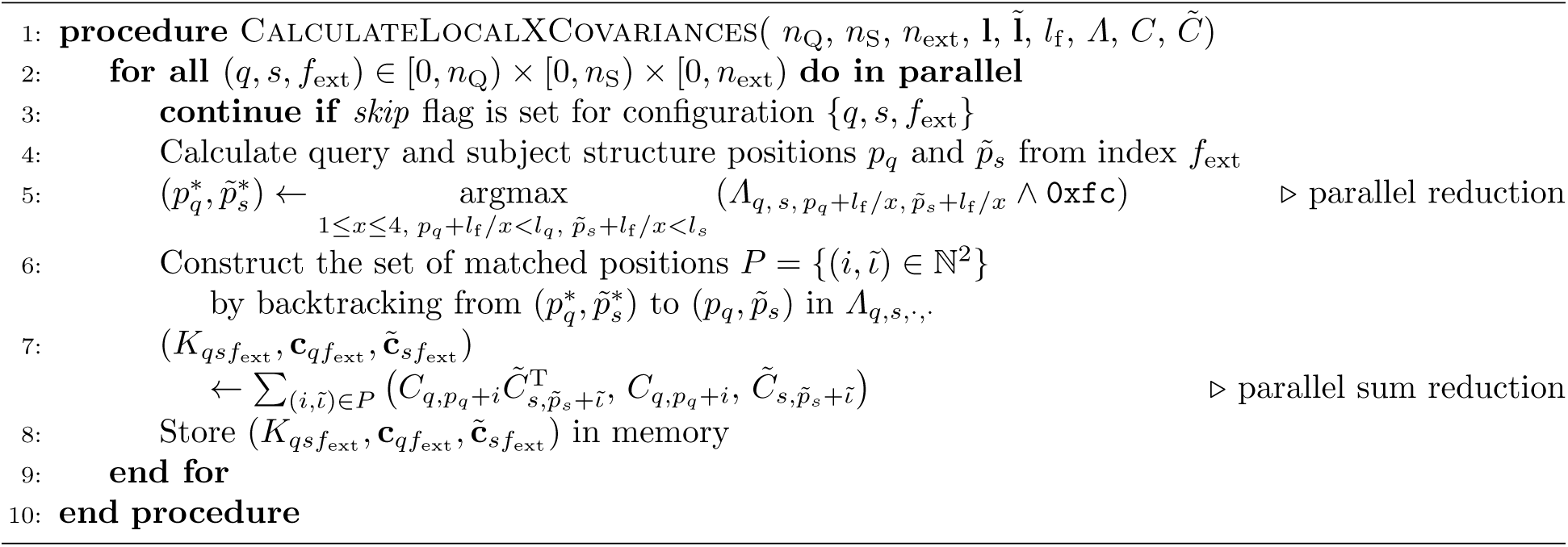
Calculate local alignment-based cross-covariance matrices.

**Algorithm 4.**
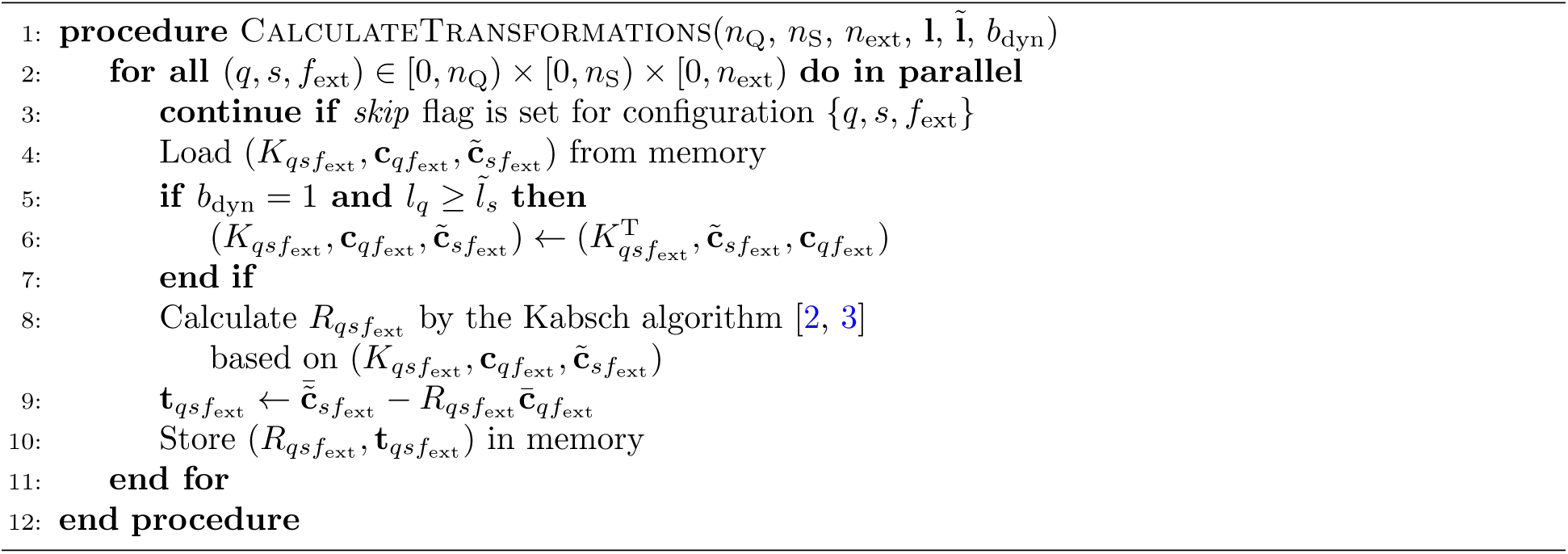
Calculate transformation matrices.

**Algorithm 5.**
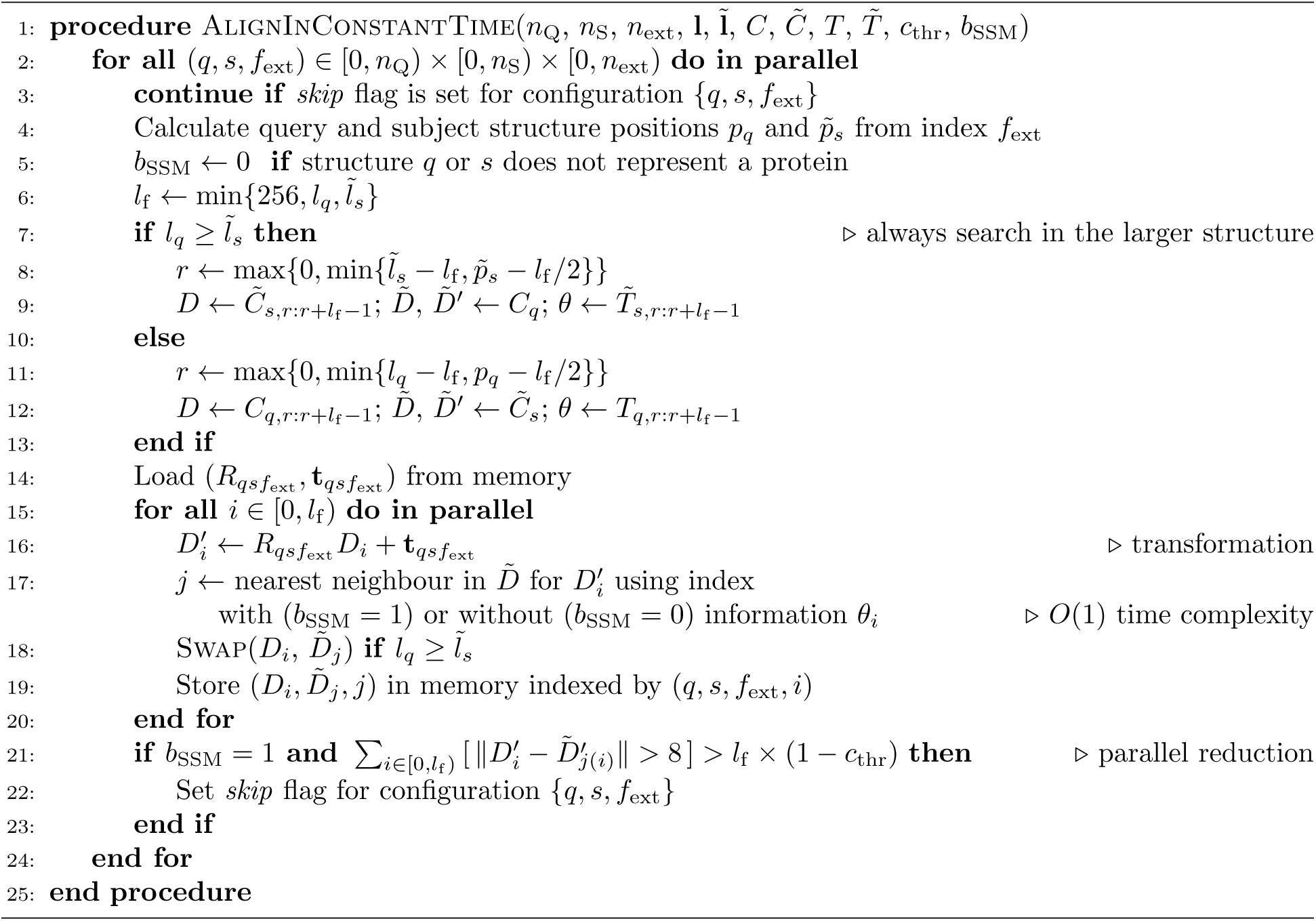
Produce pilot alignments in constant time using spatial indices.

**Algorithm 6.**
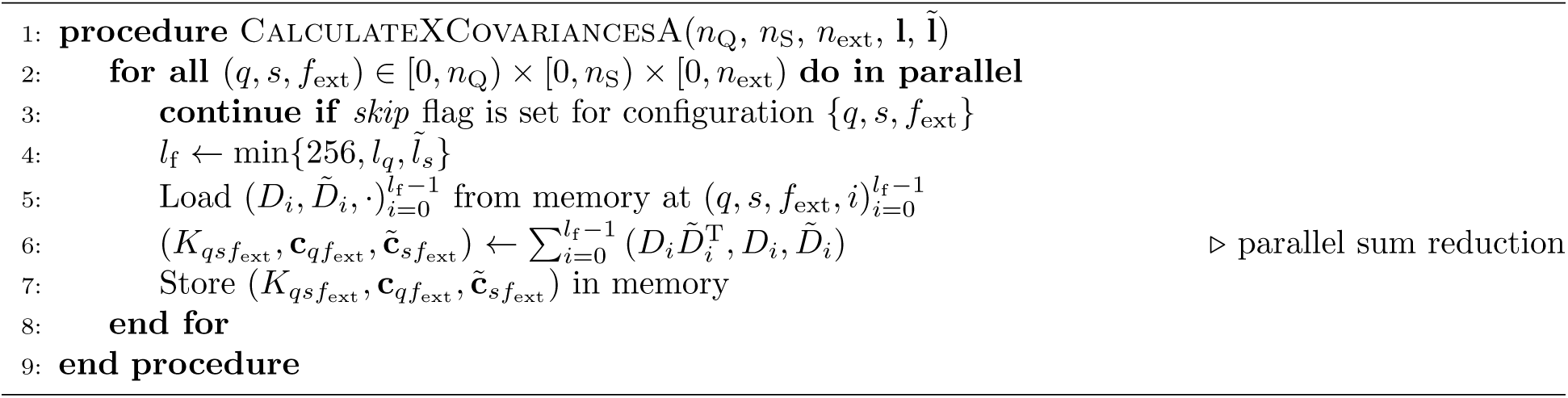
Calculate cross-covariance matrices from alignments.

**Algorithm 7.**
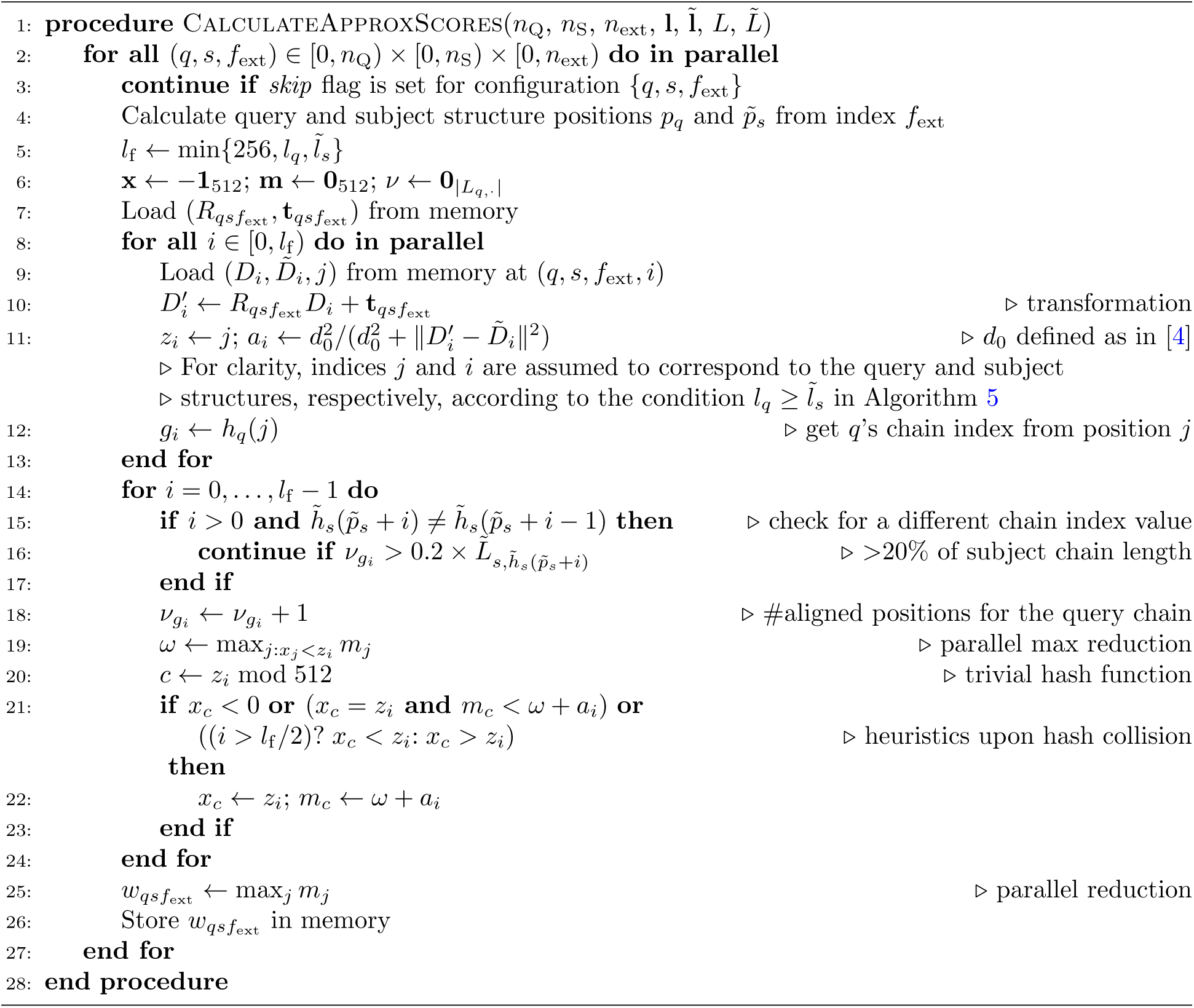
Calculate approximate sequence-order-dependent scores.

**Algorithm 8.**
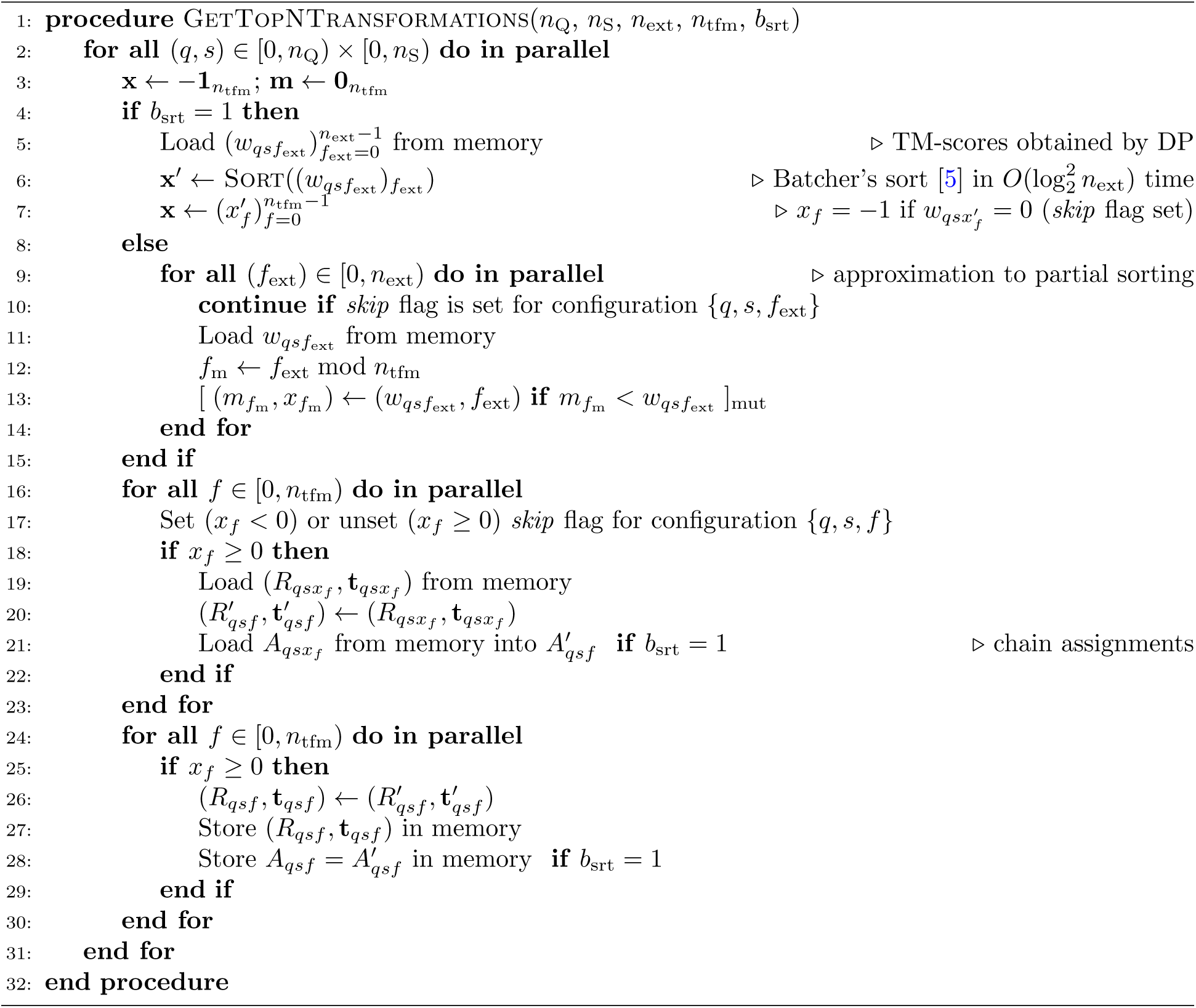
Select the top n_tfm_ transformation matrices.

**Algorithm 9.**
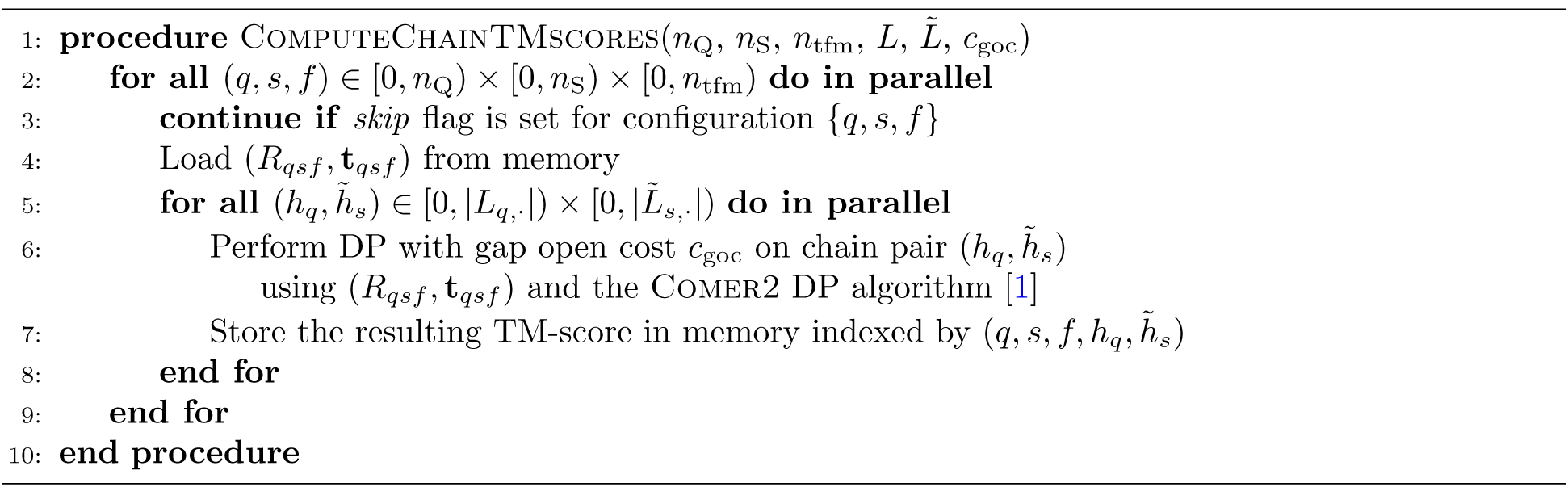
Compute chain-level TM-scores for the top n_tfm_ transformation matrices.

**Algorithm 10.**
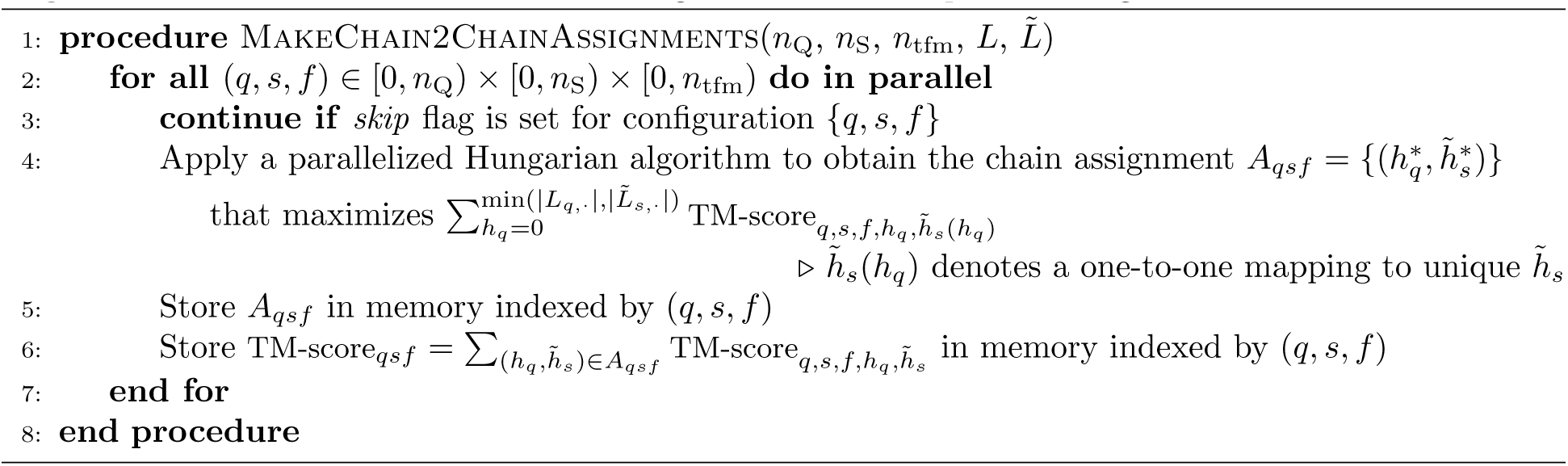
Determine chain-to-chain assignments for the top n_tfm_ configurations.

**Algorithm 11.**
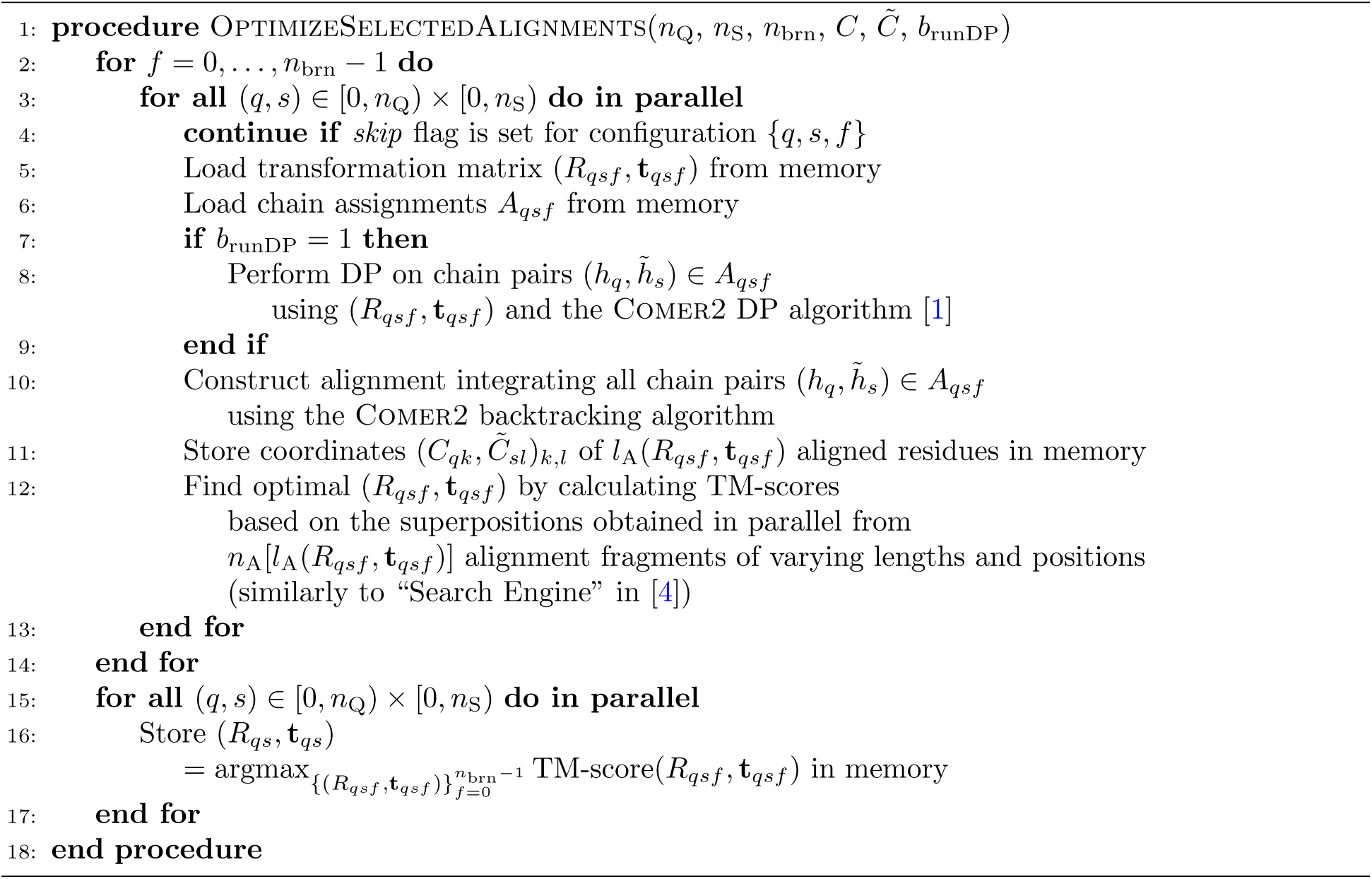
Optimize n_brn_ selected alignments.

**Algorithm 12.**
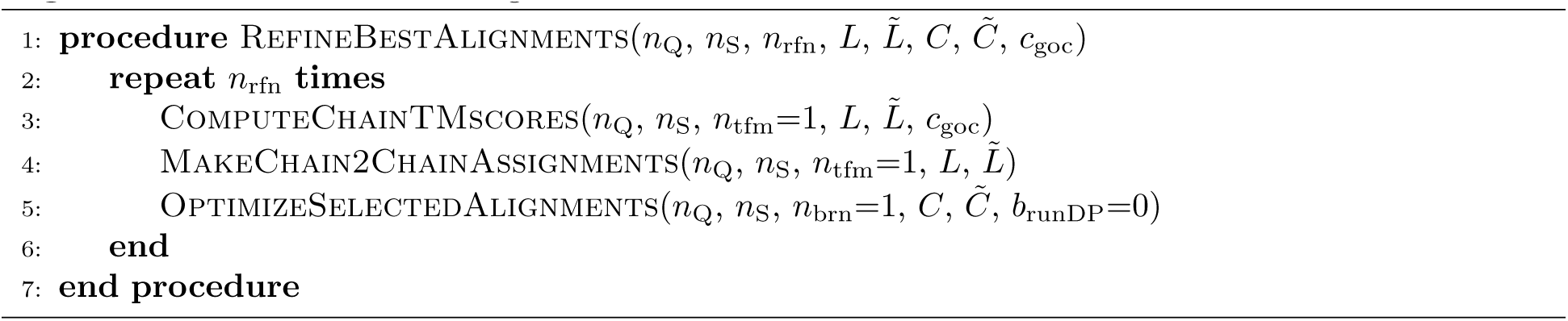
Refine the best alignments.

## Notes

### Summary of Updates

Additional datasets and benchmark studies have been added to the manuscript.

## References

[1] Harper, J. W. & Bennett, E. J. Proteome complexity and the forces that drive proteome imbalance. Nature 537, 328–338 (2016).

[2] Jumper, J. et al. Highly accurate protein structure prediction with AlphaFold. Nature 596, 583–589 (2021).

[3] Baek, M. et al. Accurate prediction of protein structures and interactions using a three-track neural network. Science 373, 871–876 (2021).

[4] Humphreys, I. R. et al. Computed structures of core eukaryotic protein complexes. Science 374, eabm4805 (2021).

[5] Baek, M. et al. Accurate prediction of proteinnucleic acid complexes using RoseTTAFoldNA. Nat. Methods 21, 117–121 (2024).

[6] Abramson, J. et al. Accurate structure prediction of biomolecular interactions with AlphaFold 3. Nature 630, 493–500 (2024).

[7] Ahnert, S. E., Marsh, J. A., Hernndez, H., Robinson, C. V. & Teichmann, S. A. Principles of assembly reveal a periodic table of protein complexes. Science 350, aaa2245 (2015).

[8] Wan, C. et al. Panorama of ancient metazoan macromolecular complexes. Nature 525, 339–344 (2015).

[9] Braberg, H. et al. Genetic interaction mapping informs integrative structure determination of protein complexes. Science 370, eaaz4910 (2020).

[10] Dey, S., Ritchie, D. & Levy, E. D. PDB-wide identification of biological assemblies from conserved quaternary structure geometry. Nat. Methods 15, 67–72 (2018).

[11] Mukherjee, S. & Zhang, Y. MM-align: a quick algorithm for aligning multiple-chain protein complex structures using iterative dynamic programming. Nucleic Acids Res. 37, e83–e83 (2009).

[12] Koike, R. & Ota, M. SCPC: a method to structurally compare protein complexes. Bioinformatics 28, 324–330 (2011).

[13] Madej, T. et al. MMDB and VAST+: tracking structural similarities between macromolecular complexes. Nucleic Acids Res. 42, D297–D303 (2013).

[14] Zhang, C., Shine, M., Pyle, A. M. & Zhang, Y. US-align: universal structure alignments of proteins, nucleic acids, and macromolecular complexes. Nat. Methods 19, 1109–1115 (2022).

[15] Kim, W. et al. Rapid and sensitive protein complex alignment with Foldseek-Multimer. Nat. Methods 22, 469–472 (2025).

[16] Minami, S., Sawada, K., Ota, M. & Chikenji, G. MICAN-SQ: a sequential protein structure alignment program that is applicable to monomers and all types of oligomers. Bioinformatics 34, 3324–3331 (2018).

[17] Sippl, M. J. & Wiederstein, M. Detection of spatial correlations in protein structures and molecular complexes. Structure 20, 718–728 (2012).

[18] Lyu, Q., Wei, H., Chen, S., Peng, Z. & Yang, J. Rapid and accurate protein structure database search using inverse folding model and contrastive learning. J. Chem. Inf. Model. 65, 13465–13477 (2025).

[19] Kåhrsträm, C. T. Type III CRISPR-Cas complexes in the spotlight. Nat. Rev. Microbiol. 11, 821 (2013).

[20] Margelevičius, M. GTalign: spatial index-driven protein structure alignment, superposition, and search. Nat. Commun. 15, 7305 (2024).

[21] Zhang, Y. & Skolnick, J. Scoring function for automated assessment of protein structure template quality. Proteins 57, 702–710 (2004).

[22] Margelevičius, M. COMER2: GPU-accelerated sensitive and specific homology searches. Bioinformatics 36, 3570–3572 (2020).

[23] Burley, S. et al. Updated resources for exploring experimentally-determined PDB structures and computed structure models at the rcsb protein data bank. Nucleic Acids Res. 53, D564–D574 (2025).

[24] Zhang, Y. & Skolnick, J. TM-align: a protein structure alignment algorithm based on the TM-score. Nucleic Acids Res. 33, 2302–2309 (2005).

[25] Zemla, A. LGA: a method for finding 3d similarities in protein structures. Nucleic Acids Res. 31, 3370–3374 (2003).

[26] Sonani, R. R., et al. Tad and toxin-coregulated pilus structures reveal unexpected diversity in bacterial type IV pili. Proc. Natl. Acad. Sci. U.S.A. 120, e2316668120 (2023).

[27] Tauriello, G. et al. ModelArchive: A deposition database for computational macromolecular structural models. J. Mol. Biol. 437, 168996 (2025).

[28] Xu, J. & Zhang, Y. How significant is a protein structure similarity with tm-score=0.5? Bioinformatics 26, 889–895 (2010).

[29] Dapkūnas, J. & Margelevičius, M. Web-based GTalign: bridging speed and accuracy in protein structure analysis. Nucleic Acids Res. 53, W291–W296 (2025).

[30] Feldman, J. & Skolnick, J. AF3Complex yields improved structural predictions of protein complexes. Bioinformatics 41, btaf432 (2025).

[31] Gong, S., Zhang, C. & Zhang, Y. RNA-align: quick and accurate alignment of RNA 3D structures based on size-independent TM-scoreRNA. Bioinformatics 35, 4459–4461 (2019).

[32] Pettersen, E. F. et al. UCSF ChimeraX: Structure visualization for researchers, educators, and developers. Protein Sci. 30, 70–82 (2021).

[33] Wickham, H. Ggplot2: Elegant graphics for data analysis 2 edn. Use R! (Springer International Publishing, Cham, Switzerland, 2016).

[34] R Core Team. R: A Language and Environment for Statistical Computing. R Foundation for Statistical Computing, Vienna, Austria (2021). URL https://www.R-project.org/.

## Bibliography

[1] Margelevičius, M. COMER2: GPU-accelerated sensitive and specific homology searches. Bioinformatics 36, 3570–3572 (2020).

[2] Kabsch, W. A solution for the best rotation to relate two sets of vectors. Acta Crystallogr. A 32, 922–923 (1976).

[3] Kabsch, W. A discussion of the solution for the best rotation to relate two sets of vectors. Acta Crystallogr. A 34, 827–828 (1978).

[4] Zhang, Y. & Skolnick, J. Scoring function for automated assessment of protein structure template quality. Proteins 57, 702–710 (2004).

[5] Batcher, K. E. Sorting networks and their applications. *Proceedings of the April 30–May 2*, *1968, Spring Joint Computer Conference* 307–314 (1968).

